# Opposing, multiplexed information in lateral and ventral orbitofrontal cortex guides sequential foraging decisions in rats

**DOI:** 10.1101/2024.08.17.608409

**Authors:** Paul J. Cunningham, A. David Redish

## Abstract

Far medial and lateral aspects of the orbitofrontal cortex (OFC) serve distinct roles in value-based decision-making, though it remains unclear how more nearby lateral and ventral aspects differ in their information-processing characteristics. The present study leveraged high-density neural recording in the lateral orbital (LO) and ventral orbital (VO) subregions using a neuroeconomic task in rats to clarify how functional heterogeneity between LO and VO participates in sequential cost-benefit foraging. LO and VO contained opposing representations of an encountered reward’s subjective benefit and its associated opportunity costs, respectively. The relative balance of these representations tracked decisions to approach or leave a reward during each stage of an encounter. Opposing representations of subjective benefit in LO and opportunity costs in VO were realized by distinct clusters of cells within each subregion across stages of a reward encounter (i.e., task state). In addition, overall activity in each subregion scaled with changes in the subjective value of rewards induced by external, economic conditions (reward scarcity) and internal, motivational factors (hunger). Collectively, these results suggest that lateral and ventral OFC encode opposing representations to guide sequential foraging decisions, but using a shared coding scheme which multiplexes information about subjective value (via overall activity) and world states (via distinct ensembles).

## Introduction

Animals must regularly decide whether to spend limited resources on a currently available reward or search for better rewards in the future^1–3^. Consider a bear as trout swim past, or a rideshare driver deciding whether to accept a current request. Such sequential foraging paradigms are key to understanding how representations of costs and benefits are integrated to guide behavior ^4–8^. The orbitofrontal cortex (OFC) participates in a wide range of processes that enable sequential cost-benefit foraging decisions^9–13^. Importantly, the OFC is anatomically and functionally heterogenous across its medial-to-lateral axis^14–22^. This raises the question of how OFC subregions differ in the representations they contain and computations they perform during sequential cost-benefit foraging decisions.

Research exploring OFC functional heterogeneity has focused primarily on comparisons between its far medial and far lateral aspects^19–24^. This work suggests that lateral OFC contributes to behaviors guided by stimuli that signal rewards^25–32^, while medial OFC participates when reward value is dynamically changing and actions require complex valuations based on unobservable information^33–38^. These functional distinctions are consistent with anatomical distinctions demonstrating that lateral aspects of the OFC are more strongly connected to sensory and motor regions implicated in stimulus-guided actions whereas medial aspects of the OFC are more strongly connected to fronto-cortical and sub-cortical regions implicated in emotional regulation, motivation, and goal-directed actions^39–42^.

An understudied but equally important functional distinction within the OFC exists between its lateral and ventral aspects. Indeed, it is common for studies to refer to the “VOLO” subregion of the OFC when experimental targeting spans both subregions, with the assumption of functional overlap between them. However, a cross-study synthesis^15^ of OFC heterogeneity suggested a distinct functional role for VO in cost-benefit decision-making. Unlike LO, which participates in stimulus-guided reward approach and anticipation, VO participates in behaviors that are temporally distant from reward and when reward acquisition requires inhibition of prepotent approach behaviors^43–48^. Thus, functional heterogeneity between lateral and ventral aspects of the OFC are just as evident as its far lateral versus far medial aspects. However, there are no studies that have directly explored how the timing and content of representations within LO and VO ensembles participate in sequential cost-benefit foraging decisions.

Restaurant Row (Figure 1a) is a neuroeconomic sequential foraging task in which subjects are given a limited amount of time to forage for consumable rewards. ^46–53^ Upon encountering a “restaurant”, subjects enter an “offer zone” (OZ) where they are informed of the delay to reward for the current offer. Accepting an offer requires entering the “wait zone” (WZ); rejecting it entails moving to the next restaurant. In the WZ, the delay counts down until reward is delivered, but subjects can quit the offer at any time by leaving the restaurant. If subjects wait out the delay in the WZ, reward is delivered and subjects enter the subsequent “linger zone” (LZ), which ends when subjects leave for the next restaurant (on their own accord). When the delay range is small (1-10s), subjects achieve their long-term consumption goals by accepting every reward. When the delay range is large (1-30s), subjects must adapt to the reward-scarce environment. Subjects initially adapt by learning to quit bad offers in the WZ, and eventually learn to reject bad offers in the OZ rather than entering the restaurant and wasting time waiting before quitting^56^. Thus, sequential cost-benefit decisions on Restaurant Row are defined by an initial valuation in the OZ followed by a subsequent reevaluation in the WZ. Initial evaluations and subsequent reevaluations are governed by distinct processes within the frontal cortex, striatum, and hippocampus^49,51,54,55^. Restaurant Row therefore provides an ideal platform to study the diverse computational roles of lateral and ventral OFC in sequential foraging.

**Figure 1.**
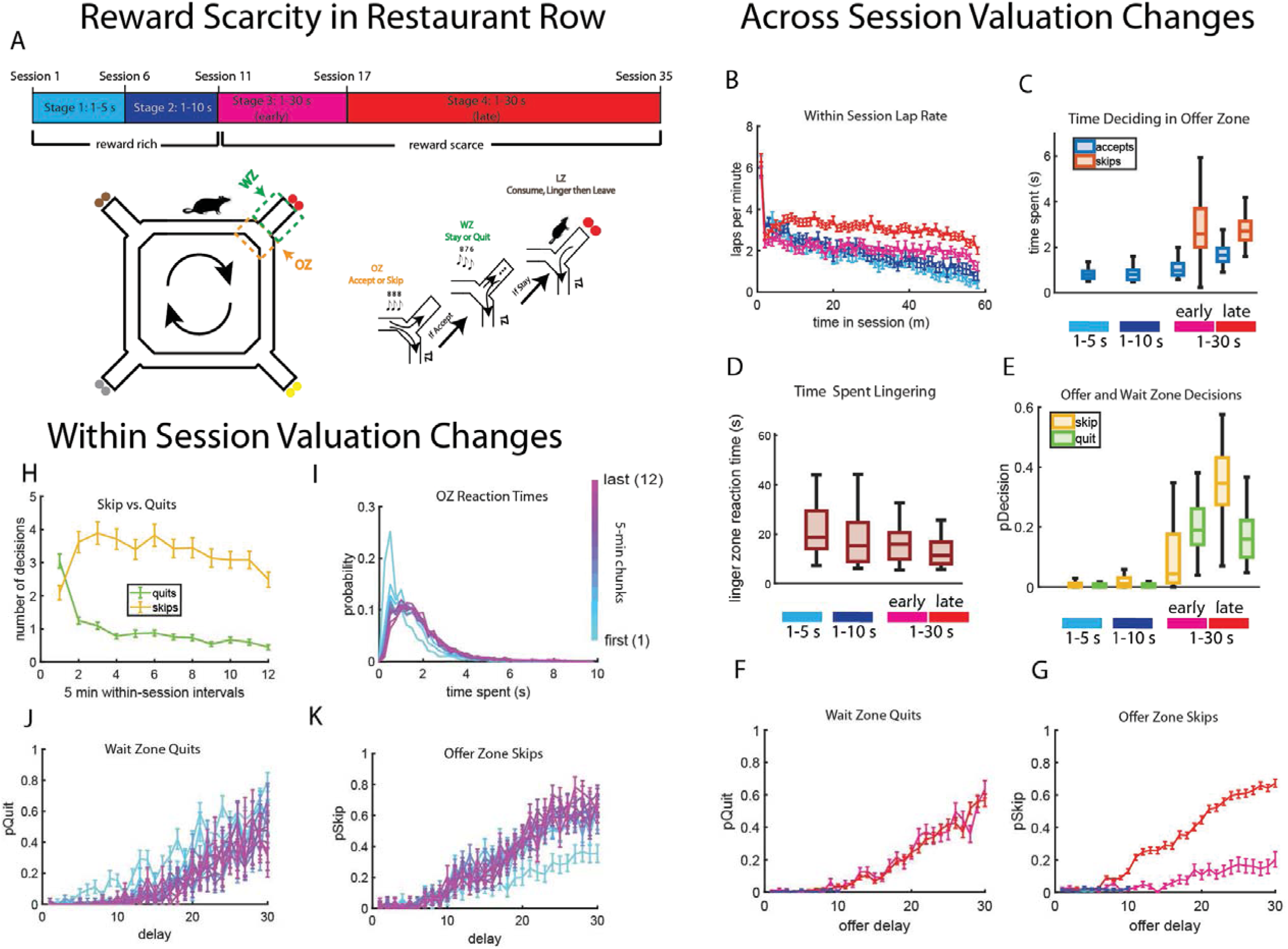
Behavioral adaptations to a reward scarce environment in Restaurant Row. (A) Restaurant Row is a maze containing four restaurants offering distinct foods following a delay. Recording sessions were categorized into four stages based on reward availability within 1 hour foraging sessions. (B) Mean (±SEM) laps per minute averaged across sessions in each of the four training stages. (C) Time spent making decisions in the offer zone across the four training stages. (D) Time spent lingering after reward across the four training stages. (E) Probability of skipping in the OZ (orange) and quitting in the WZ (green) across the four stages of training. (F) Mean (±SEM) probably of quitting across delays across training stages. (G) Mean (±SEM) probability of skipping across delays across training stages. (H) Mean (±SEM) number of skips and quits across successive 5-min session chunks. (I) Probability density functions of time spent making decisions in the OZ across 5-min session chunks. (J) Mean (±SEM) probability of quitting as a function of delay across 5-min session chunks. (K) Mean (±SEM) probability of skipping as a function of delay across 5-min session chunks.

The present study leveraged high-density neural ensemble recordings of lateral orbital (LO) and ventral orbital (VO) cortices while rats navigated Restaurant Row. We discovered distinct neural dynamics between OFC subregions. LO reflected decisions to approach and wait for reward by encoding the current reward’s subjective benefits. In contrast, VO reflected decisions to leave the current restaurant and search for reward elsewhere by representing the opportunity costs incurred by waiting for the current reward. Distinct ensembles in LO and VO participated in these representations along the sequence of events (task stages) that defined a reward encounter. These decision-relevant representations were sensitive to both external, economic factors (i.e., reward scarcity) and internal, motivational factors (i.e., hunger). Thus, while LO and VO contained opposing representational content, they both multiplexed information about task state and subjective value via a common coding scheme (i.e., distinct ensembles representing task state with overall activity representing subjective value).

## Results

### Behavior in Restaurant Row adapted to economic and motivational factors

Rats foraged for their daily food ration during 60 min sessions on Restaurant Row. Each restaurant offered a distinctly flavored pellet, and flavor assignment to restaurant locations was fixed throughout the experiment. Upon encountering a restaurant, the delay to reward for that encounter was randomly drawn from a range. Across blocks of sessions, this delay range increased, making rewards more scarce as training progressed (with session blocks referred to as “stages”; Figure 1A). During stage 1, rewards were delivered following a 1-5 s delay. During stage 2, rewards were delivered following a 1-10 s delay. Following 5 sessions at stages 1 and 2 each, the delay range increased to 1-30 s, which defined the reward scarce environment and lasted 25 sessions. Based on behavioral adaptations described below, this reward scarce environment was broken up into an “early” and “late” stage. Stage 3 (early) refers to the first 7 session of the reward scarce environment. During this stage, rats were adjusting to the new, reward-scarce environment. Stage 4 (late) refers to the remaining 18 sessions of the reward scarce environment and reflects the period after behavioral adaptations stabilized. Because session duration was fixed, increasing the delay range imposed greater opportunity costs - time spent waiting for the current reward was time lost that could have been spent searching for more profitable rewards. Thus, rewards with longer delays were associated with greater opportunity costs. We first explored how behavior changed across training stages, reflecting adaptations to the reward scare environment. Because we did not find consistent flavor preferences (S1A-G), all behavioral analyses were performed on data merged across restaurants.

Prior to the reward-scarce environment, rats would forage at a constant pace for the first half of the session but took breaks near the end of the session, as reflected in decreasing lap rate as session time progressed (Figure 1B; ANOVA – main effect of session time: F(57,11477) = 34.59, p< 0.0001). This pattern carried over into the early portion of the reward-scarce environment but was eventually replaced by a relatively constant rate of lap completions throughout the session during the final stage of the reward scarce environment (Figure 1B; ANOVA – main effect of stage: F(4,11477) = 551.53, p< 0.0001; stage×session-time interaction: F(171,11477)=2.33, p<0.0001). The more consistent rate of lap completion that emerged later in the reward-scarce environment was accompanied by an increase in time spent making decisions in the offer zone (OZ; Figure 1C; ANOVA – main effect of stage: F(3,251)=117.03, p=1.9×10^-47^) and a decrease in time spent lingering after reward (Figure 1D; ANOVA – main effect of stage: F(3,251)=17.91, p=1.5×10^-10^). These results suggest that introduction to the reward-scarce environment changed how rats spent their limited foraging time. When rewards were abundant, rats would front-load their foraging bouts at the start of each session and take breaks thereafter. When rewards were scarce, this strategy resulted in a reduction in food intake (S1H) and therefore encouraged rats to make more careful decisions based on reward delay (S1I).

In addition to reallocating temporal resources, rats changed their decision-making strategies when rewards became scarce. Prior to the reward-scarce environment, rats accepted and earned every offer (Figure 1E). Upon introduction to the reward-scarce environment, rats began quitting offers in the wait zone (Figure 1E). Importantly, quits were more likely for longer delays (Figure 1F; ANOVA – main effect of delay: F(1,4137)=2137.01, p<.0001). Sensitivity of quits to reward delay was constant across training stages (Figure 1F; ANOVA - main no main effect of between stages 3 and 4: F(1,5082)=0.02, p=0.89; no delays×stage interaction: F(1,5082)=0.22, p=0.64).

Within approximately 1 week of exposure to the reward-scarce environment, skips became more frequent than quits (Figure 1E; ANOVA – stage×decision interaction: F(10,3784)=12.6, p<.0001), suggesting that rats learned to avoid entering a restaurant rather than entering then leaving the restaurant before reward was delivered. Like quits, the probability of skipping increased with delay (Figure 1G; ANOVA – main effect of delay: F(1,4279)=2855.01, p<.0001). Unlike quits, however, the sensitivity of skips to delay increased with experience in the reward-scarce environment (ANOVA - main effect of delay: F(1,5242)=1729.07, p<.0001; main effect between stages 3 and 4: F(1,5242)=23.92, p<.0001; significant delay×stage interaction, F(1,5242)=635.54, p<.0001).

Time spent in the OZ increased with experience in the reward-scarce environment, with skips taking longer than accepts (Figure 1C; Mann-Whitney U Test - z=-55.31, p<.001). Interestingly, short-delay offers were accepted more quickly than long-delay offers, whereas long-delay offers were skipped more quickly than short-delay offers (Figure S1 J-K; ANOVA - main effect of decision: F(1,6366)=332.81, p<.0001; delay×decision interaction: F(1,6366)=12.15, p<.0001), suggesting a fast-acting accept process overridden by a slower skip process that were both sensitive to delay.

Although rats made session-wide adaptations to the global economic climate of the task (defined by reward delays across stages), there was an initial period at the start of each session which reflected a fundamentally different decision-making strategy (see the initial burst of laps at the start of each session in Figure 1B). Rats were more likely to accept and then quit, and less likely to skip, during the first 5 min of each session than any subsequent 5-min epoch within the session (Figure 1H; ANOVA – main effect of session chunk: F(10,3784)=2.86, p=0.001; main effect of decision: F(1,3784)=613.49, p<0.0001; chunk×decision interaction: F(10,3784)=12.6, p<0.0001). Sensitivity of quits to delay was relatively constant throughout the session (Figure 1J; ANOVA – no main effect of chunk on 50% choice threshold: F(10,478)=0.57, p=0.83). However, skip decisions were less sensitive to offer delay during this first 5 min epoch (Figure 1K; ANOVA – main effect of chunk on 50% choice threshold: F(10,715)=3.73, p=6.7×10^-5^), indicating increased willingness to accept rewards with longer delays. Insensitivity to offer delay was accompanied by less time spent making decisions in the OZ during the initial 5 min (Figure 1I; ANOVA – main effect of chunk: F(10,21664)=162.28, p=8.1×10^-90^). Thus, the start of each session was characterized by a brief period in which rats overvalued reward.

Collectively, these results suggest that the subjective value of rewards in Restaurant Row depends on 1) external, economic conditions determined by reward scarcity, and 2) internal, motivational factors determined by regulatory processes related to hunger. Such changes in subjective value guide behavioral adaptations both between sessions (economic) and within session (motivation). The question is, how do neural ensembles within lateral (LO) and ventral (VO) aspects of the OFC encode decision-relevant information to support adaptations to economic and motivational factors? To address this, we explored how neural dynamics in LO and VO ensembles (see S2 for histology) during key decision epochs in Restaurant Row were affected by global and local economic conditions.

### Representations of subjective value and opportunity costs in LO and VO adapted to economic conditions

OFC plays a critical role in integrating multidimensional information to make cost-benefit decisions^20,57–62^. OZ decisions in Restaurant Row depend on sensory information about reward delay, learned relations between tone pitch and delay, spatial location, and knowledge about reward-restaurant associations to infer the value of current offers relative to future opportunities. It is unclear how LO and VO participate in such multidimensional cost-benefit decisions. To this end, we compared neural dynamics throughout the OZ and found distinct activity profiles between subregions at OZ entry (Mixed Effects Model - main effect of region: F(1,8402)=361.62, p=5.0×10^-80^; main effect of time: F(1,8402)=60.87, p=6.2×10^-16^; significant time x region interaction: F(1,8402)=13.98, p=1.7×10^-7^) and at OZ exit (Mixed Effects Model - main effect of region: F(1,8382)=317.9, p=3.8×10^-71^; main effect of time: F(1, 8382)=5.72, p=0.001; significant time x region interaction: F(1, 8382)=0.93, p=0.03). To clarify these differences, we first examined representational content and timing of ensemble activity in LO and VO after rats adapted behavior to the reward scarce environment (stage 4).

LO ensembles exhibited a burst of activity when rats entered the OZ and the tone signaling reward delay was presented (Figure 2A). This burst began prior to tone onset and did not contain information about delay (Mixed Effects Model – no main effect of delay: F(1,3417)=0.003, p=0.1). Interestingly, after the initial burst, LO activity remained elevated during later periods of the OZ only when offer delay was relatively short (Figure 2A-B). The timing of this second neural signature in LO corresponded roughly to the time at which rats made decisions in the OZ (i.e., after the first tone was presented and they had time to process information about the current offer). Cell-averaged LO tuning curves to reward delay during this putative decision period revealed a non-linear relation between neural activity and reward delay reminiscent of functions that relate subjective value to delay (Figure 2E; Mixed Effects

**Figure 2.**
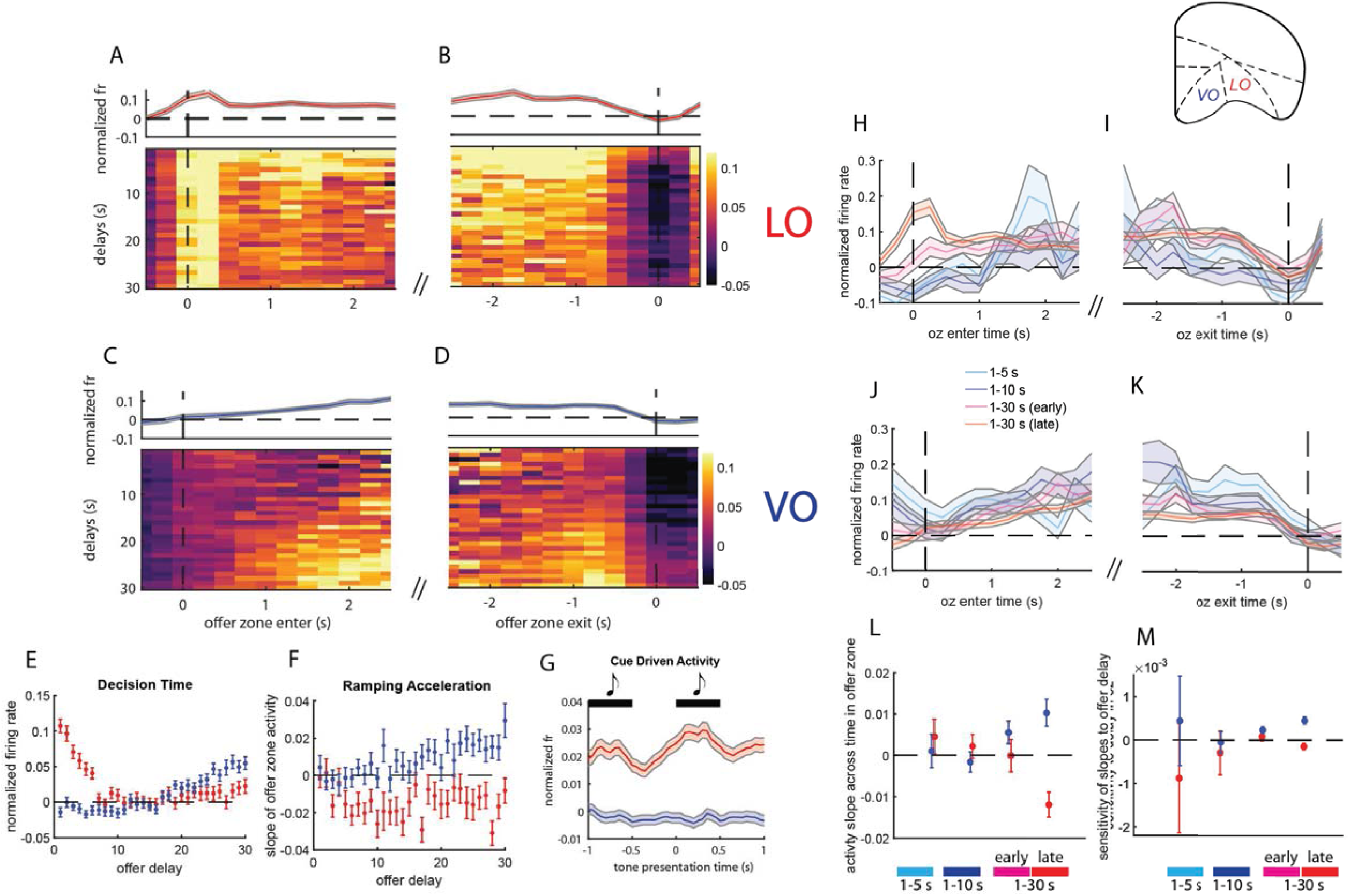
Opposing representations in LO and VO adapted to economic conditions in the OZ. (A-B) Mean neural activity in LO as a function of delay at OZ entry (A) and exit (B). Top panel shows mean (±SEM) activity averaged across delays. (C-D) Same as A-B but for VO activity. (E) Mean (±SEM) tuning curves of LO (red) and VO (blue) to offer delay during the decision period in the OZ. (F) Mean (±SEM) sensitivity of ramping activity to offer delay for LO and VO. (G) Mean (±SEM) activity in LO and VO oriented to tone presentations in the OZ. (H-I) Mean (±SEM) neural activity in LO at OZ entry (H) and exit (I) across the four training stages. (J-K) Same as H-I but for VO activity. (L) Mean (±SEM) degree of ramping activity in the OZ across the four training stages. (M) Mean (±SEM) sensitivity of ramping activity to offer delay in the in OZ across the four training stages.

Model - main effect of delay: F(1,3535)=98.41, p=7.4×10^-24^). Finally, consistent with the notion that LO encoding is linked to environmental cues, periods of elevated LO activity in the OZ was temporally aligned to tone presentation, with rhythmic activity reaching its peak moments before the tone turned off (Figure 2G), VO activity steadily increased with time spent in the OZ, ramping more steeply for offers with longer delays (Figure 2C-D). To quantify how VO ramping depended on the interaction between offer delay and time spent in the OZ, we measured the slope of neural activity as a function of time spent for each delay. Steepness of the activity ramp, indicating acceleration in neural activity, increased with offer delay (Figure 2F; Mixed Effects Model – main effect of delay: F(1,2848)=45.29, p=2.9×10^-12^). Further, cell-averaged VO tuning curves to delay during the putative OZ decision period revealed a positive relation between activity and delay (Figure 2E; Mixed Effects Model – main effect of delay: F(1,2846)=278.15, p=1.6×10^-60^). Finally, unlike LO, VO activity in the OZ was unaffected by tone presentations (Figure 2G).

Because offer delay is critical to infer subjective value, delay-dependent dynamics in LO and VO might reflect the subjective value of an offer rather than delay *per se*. The reward-scarce environment induced changes in the subjective value of offers that encouraged decisions informed by costs and benefits. Thus, if delay-dependent activity in LO and VO reflected subjective valuations, activity in each subregion should be affected by introduction and adaptation to the reward scarce environment. To address this possibility, we compared neural dynamics in the OZ across training stages (Figure 1A).

The burst of LO activity at OZ entry emerged only after rats adapted to the reward-scarce environment (i.e., stage 4; Figure 2H; Mixed Effects Model – main effect of stage: F(3,183)=10.45, p=5.5×10^-7^; post-hoc comparison between stage 4 and stages 1-3: t(183)=3.1, p=0.002; post-hoc comparison between stage 3 and stages 1,2, and 4: t(183)=0.14, p=0.88). This suggests that LO activity marked cue onset only after that cue acquired behavioral relevance.

Further, while VO activity ramped to some degree throughout training Figure 2J-K), this ramping was not affected by offer delay until after the reward-scarce environment was introduced (Figure 2L-M; Mixed Effects Model – main effect of stage: F(2,212)=3.65, p=0.001). Importantly, delay-dependent VO ramping emerged prior to behavioral adaptations in the reward scarce environment and therefore preceded changes in cost-benefit decisions (Figure 2L-M; Mixed Effects Model – main effect between stages 2 and 3: t(212)=2.5, p=0.01; main effect between stages 2 and 4: t(212)=2.3, p=0.01). Thus, cue-driven activity linked to short delays in LO, along with ramping activity linked to long delays in VO, did not reflect absolute delay but instead reflected subjective valuations of offers that tracked changes in global economic conditions.

### Opposing representations in LO and VO participated in offer zone decisions and adapted to motivational conditions

Sequential foraging decisions are made by comparing the value of currently encountered rewards with the opportunity costs associated with pursuing that reward^63–66^. In Restaurant Row, the current reward’s value is determined by its delay while the opportunity costs are determined by the average expected delay to reward in the environment. Within this context, LO activity represented the current offer’s subjective value through heightened activity for good offers early in the OZ. In contrast, VO activity represented the opportunity costs lost by waiting for the current reward through gradual ramping activity that appeared later in the OZ. The content and timing of these opposing representations suggest that the relative activity of LO to VO might serve as an integrated signal of offer value that can be used for accept-skip decisions.

Consistent with this hypothesis, there were decision-dependent differences between LO and VO activity that emerged after OZ entry (S3A-D: Mixed Effects Model – main effect of region: F(1,15692)=401.9, p=0.01; no main effect of decision: F(1,15692)=0.01, p=0.07) and lasted until OZ exit (Mixed Effects Model – main effect of region: F(1,15549)=10.71, p=8.9×10^-6^; main effect of decision: F(1, 15549)=87.93, p=6.4×10^-22^). Early in the OZ, LO activity exceeded VO activity, regardless of which decision was made (Figure 3A, 3C, and 3E; Mann-Whitney U Test - comparing LO-to-VO activity at OZ entry; accepts: z = 3.8, p=.3×10^-4^; skips: z=2.2, p=0.02). If the offer was accepted, increased activity in LO relative to VO was maintained throughout the rest of the OZ (Figure 3B and 3F). If the offer was skipped, VO activity overtook LO activity towards the end of the OZ (Figure 3D and 3F; Mann-Whitney U Test: comparing LO-to-VO activity during skips at zone exit: z=-2.4, p=0.02). As a result, accept decisions were associated with a greater proportion of time-spent in the OZ with LO activity exceeding VO activity compared to skips (S3E-I). These results are consistent with the notion that LO ensembles encoded information about the current offer’s value to support fast accept decisions whereas VO ensembles encoded information about future opportunity costs to support slower skip decisions. Information about the current offer’s subjective value in LO coupled with its opportunity costs in VO can therefore be interpreted as an integrated representation of value used to make decisions. Importantly, however, these opposing representations were not in direct competition in real time, indicated by a lack of moment-by-moment correlation in LO and VO ensemble activity (S3J-K). Thus, opposing representations in LO and VO were likely unrelated to the momentary accumulation of evidence for or against a particular decision, but were instead related to providing information needed for such an accumulation process realized elsewhere.

**Figure 3.**
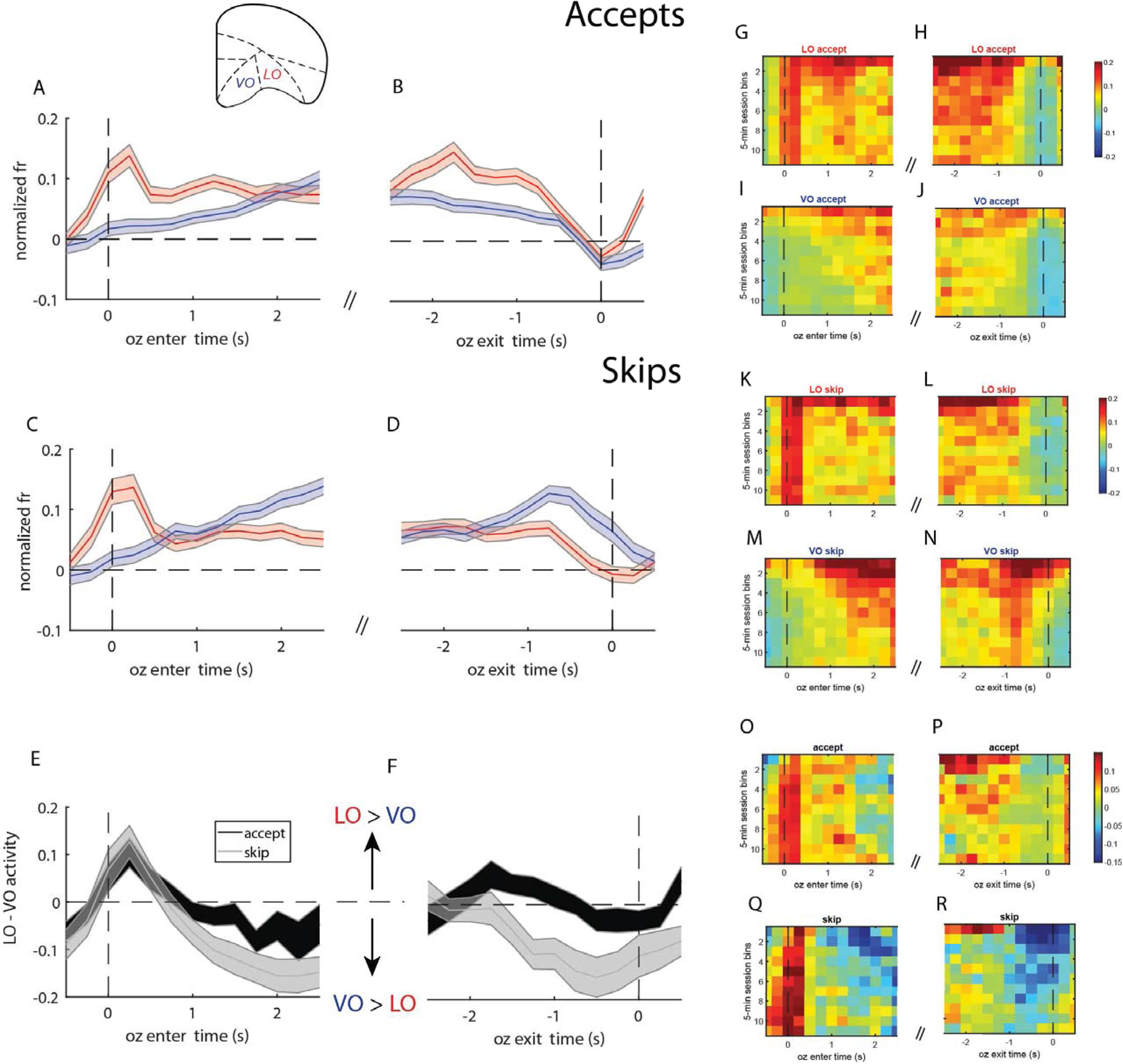
Decision-related activity in LO and VO adapted to motivational factors. (A-B) Mean (±SEM) activity in LO (red) and VO (blue) at OZ entry (A) and exit (B) when offers were accepted. (C-D) Same is in A-B but for offers that were skipped. (E-F) Mean (±SEM) relative LO-to-VO activity expressed as the momentary difference between LO and VO ensembles at OZ entry (E) and exit (F). The darker line shows relative LO-to-VO activity during accepts while the lighter line shows the same for skips. (G-H) Mean LO activity at OZ entry (G) and exit (H) during accepts, broken up into successive 5-min chunks across session time (y axis). (I-J) Same as in G-H but for VO activity. (K-L) Mean LO activity at OZ entry (K) and exit (L) during skips, broken up into successive 5-min chunks. (M-N) Same as in K-L but for VO activity. (O-P). Mean difference in LO- to-VO activity during accepts at OZ entry (O) and exit (P) broken up into 5-min chunks. (Q-R) Same as in O-P but during skip encounters.

While behavior adapted on a global scale to the reward-scarce environment, the start of each session was characterized by a brief period of overvaluation of reward that persisted throughout the reward-scarce environment. Reward overvaluation during this period was reflected in fast, impulsive accept decisions that were independent of reward delay. If LO and VO activity were indeed critical to make accept and skip decisions, their activity profiles should be distinct during this initial period of reward overvaluation. Specifically, if LO activity reflected the subjective value of current rewards, LO activity should be elevated during the first 5 min of the session. If VO activity reflected the subjective opportunity costs associated with future rewards, VO activity should also be elevated during the first 5 min of the session. To address this possibility, we compared neural activity at OZ entry and exit between accept and skip decisions across successive 5-min chunks of the session.

LO activity in the OZ was highest during the first 5 min of the session and decreased as session time progressed (Figure 3G-H and 3K-L; Mixed Effects Model – main effect of session chunk: F(10,2309)=4.95, p=3.9×10^-7^). This was true during both accepts (Figure 3G-H) and skips (Figure 3K-L; Mixed Effects Model – no session-chunk×decision interaction: F(10,2299)=0.74, p=0.68). Interestingly, however, the LO burst at OZ entry was consistent throughout the entire session (Figure 3G and 3K; Mixed Effects Model – no main effect of chunk at OZ entry: F(10,2311)=0.34, p=0.96), suggesting that this feature of LO was independent of subjective value. VO activity was also highest during the first 5 min of the session and decreased as session time progressed (Figure 3I-J and 3M-N; Mixed Effects Model – main effect of chunk: F(10,1774)=7.7, p=3.5×10^-12^). The decreasing trend of VO activity in the OZ across session time occurred during both accepts (Figure 3I-J) and skips (Figure 3M-N; Mixed Effects Model: no session-chunk×decision interaction: F(10,1764)=0.95, p=0.47).

Because the first 5 min of each session was characterized by increased accept decisions, we predicted that LO activity would be elevated to a greater degree than VO activity during this period. Surprisingly, while activity between LO and VO differed between decisions, their relative balance across session time was consistent (Figure 3O-P), and this stability in relative activity across session time was true during both accepts (Figure 3O-P) and skips (Figure 3Q-R; Mixed Effects Model – no main effect of chunk accepts: F(10,874)=0.46, p=0.91; Mixed Effects Model – no main effect of chunk during skips: F(10,593)=0.42, p=0.93). Thus, while activity in LO and VO scaled with motivation-induced changes in subjective value, their relative balance remained constant.

### LO and VO differentially encoded time remaining and time spent in the wait zone

The wait zone (WZ) is a dynamically changing epoch in which temporal progression to reward was signaled by a descending sequence of tones, providing momentary information about time remaining. Such future-oriented information is distinct from past-oriented information about how much time has already elapsed in the WZ. This raises the question of the extent to which LO and VO ensembles encoded information about time spent and time remaining. We addressed this question by assessing the sensitivity of neural activity to temporal parameters of reward delivery in the WZ after behavior adapted to the reward scarce environment (stage 4).

Given the structure of the WZ countdown, each offer of a given initial delay was associated with a unique trajectory through the two-dimensional space of time remaining and time spent. To visualize this, we plotted cell-averaged tuning curves as a function of time remaining and time spent across each initial offer delay (Figure 4A-B). Both LO and VO ensembles revealed distinct activity profiles to temporal parameters in the WZ (Mixed Effects Model – main effect of region: F(1,85682)=16.07, p=4.7×10^-6^; region x time-spent×time-remaining interaction: F(1,85682)=10.80, p=6.7×10^-5^). LO ensembles represented both time spent and time remaining (Figure 4A, 4C, and 4D; Mixed Effects Model – main effect of time-remaining: F(1,47493)=42.55, p=6.3×10^-12^; main effect of time-spent: F(1,47493)=114.22, p=1.3×10^-27^). Representations of time spent were driven primarily by a burst of activity soon after entering the WZ (see next section). Sensitivity of LO activity to time remaining was driven primarily by cue presentations, as evidenced by rhythmic LO bursting time-locked to tone presentations (Figure 4E). In contrast, VO ensembles represented time remaining but not time spent (Figure 4B, 4C, and 4D; Mixed Effects Model – main effect of time-remaining: F(1,38189)=63.28; no main effect of time-spent: F(1,38139)=0.03, p=0.1). Similar to LO, VO representations of time remaining were defined by rhythmic activity in which normalized firing rate peaked at the end of each tone presentation (Figure 4E). However, the degree to which VO ensembles represented time remaining was weaker than in LO (Figure 4D; Mixed Effects Model – region×time-remaining interaction: F(1,85682)=0.44, p=0.03).

**Figure 4.**
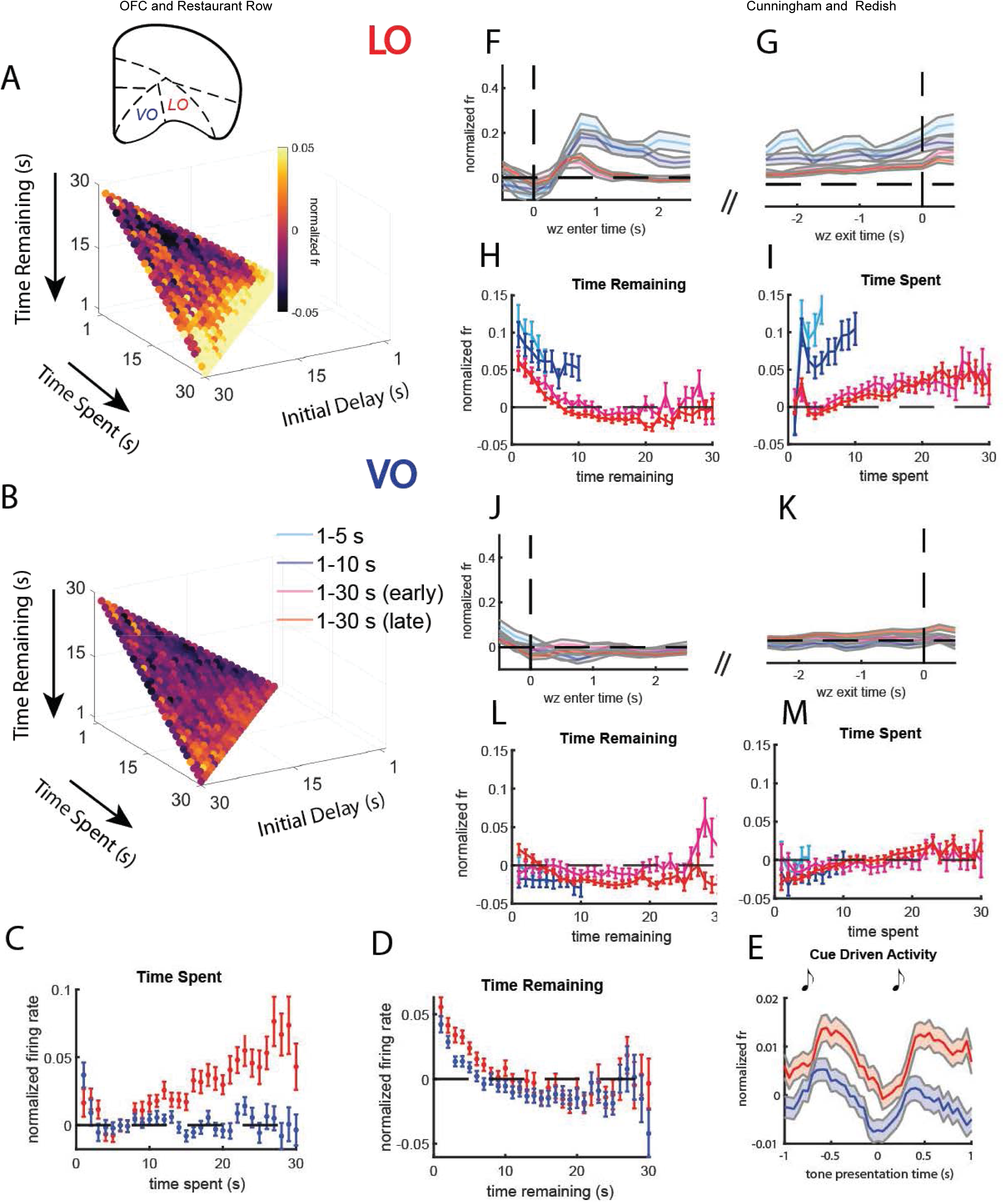
Representations of time spent and time remaining in the WZ adapted to economic conditions. (A-B) Mean tuning curves of LO (A) and VO (B) ensembles to time spent and time remaining, broken up by the initial delay that defines a trajectory of time-spent × time-remaining. (C-D) Mean (±SEM) tuning curves to time spent (C) and time remaining (D) marginalized across all other temporal variables in the WZ for LO (red) and VO (blue) ensembles. (E) Mean (±SEM) neural activity in LO and VO aligned to tone presentations in the WZ. (F-G). Mean (±SEM) LO activity at OZ entry (J) and exit (K) across the four training stages. (H-I) Mean (±SEM) tuning curves of LO activity to time remaining (H) and time spent (I) during each training stage. (J-K). Same as in F-G but for VO activity. (L-M) Same as in H-I but for VO activity.

While these results highlight distinct representations and neural dynamics in the WZ between LO and VO, it is unclear how ensembles in each subregion adapted to economic conditions. To address this question, we first compared neural activity at WZ entry and exit across the four stages of training. Surprisingly, the burst of LO activity at WZ entry was more prominent prior to the reward-scarce environment and decreased once the reward scarce environment was introduced (Figure 4F; Mixed Effects Model – main effect of stage: F(3,183)=2.22, p=0.02; follow-up comparison between stage 4 and stages 1-3: t(183)=-2.3, p=0.02; follow-up comparison between stage 3 and stages 1,2, and 4: t(183)=-2.3, p=0.02). LO activity in the moments leading up to WZ exit also decreased during reward scarce environment (Figure 4G; Mixed Effects Model – main effect of stage: F(3,183)=4.45, p=0.001; follow-up comparison between stage 4 and stages 1-3: t(183)=-2.3, p=0.02). Decreased LO activity at WZ entry and exit occurred immediately after encountering the reward scarce environment and therefore reflected the overall economic conditions rather than behavioral adaptations. In contrast, VO activity in the WZ was consistent across all stages of training (Figure 4J-K).

To explore how representations of temporal parameters in the WZ were affected by economic conditions, we compared cell-averaged tuning curves to time remaining and time spent in both LO and VO ensembles. LO activity was elevated across all values of time spent and time remaining prior to the reward scarce environment (Figure 5H-I; Mixed Effects Model – main effect of stage: F(1,4271)=29.19, p=6.9×10^-8^). However, sensitivity of LO activity to both time remaining and time spent was observed both before and after introduction to the reward-scarce environment (Mixed Effects Model – main effect of delay: F(1,4271)=14.5, p=0.0001 but no stage x delay interaction: F(1,4271)=2.21, p=0.13). Thus, while global economic conditions affected the overall level of LO activity in the WZ, they did not affect LO’s encoding information about temporal parameters. In contrast, in VO neither overall activity nor sensitivity of ensemble activity to time remaining or time spent were affected by the reward-scarce environment (Figure 4L-M; Mixed Effects Model – main effect of delay: F(1,3249)=27.54, p=1.6×10^-7^; no main effect of stage: F(1,3249)=0.82, p=0.36).

**Figure 5.**
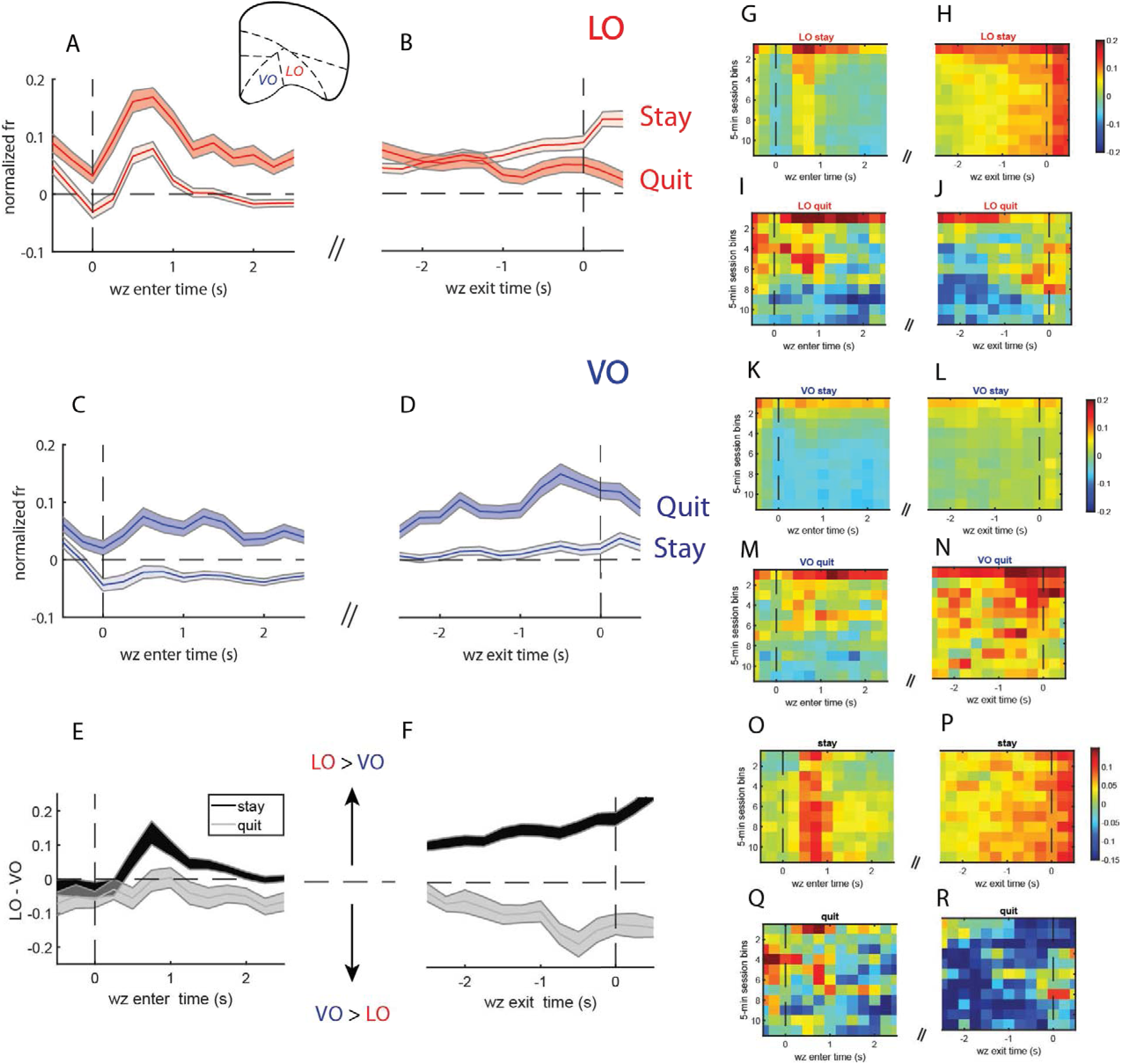
Decision-related activity in LO and VO adapted to motivational factors in the WZ. (A-B) Mean (±SEM) LO activity at WZ entry (A) and exit (B) during stay (lighter) and quit (darker) encounters. (C-D) Same as in A-B but for VO activity. (E-F) Mean (±SEM) relative LO-to-VO activity at WZ entry (E) and exit (F) during stay encounters (darker) and quit encounters (lighter). (G-H) Mean LO activity at OZ entry (G) and exit (H) during stay encounters broken up into 5-min session chunks. (I-J) Mean LO activity at WZ entry (I) and exit (J) during quit encounters broken up into 5-min session chunks. (K-L) same as in G-H but for VO activity. (M-N) same as in I-J but for VO activity. (O-P) Mean relative LO-to-VO activity during stay encounters at WZ entry (O) and exit (P) broken up into 5-min session chunks. (Q-R). Same as in O-P but during quit encounters.

### LO and VO differentially participated in reevalutive quit decisions that adapted to motivational factors

Representations of time remaining and time spent are critical for making decisions about whether to continue waiting for reward or quit in the WZ. Thus, we expected that LO and VO differentiated stays from quits. To address this, we compared neural activity in LO and VO between earn and quit decisions at WZ entry and exit. There were significant decision-dependent differences between LO and VO at both WZ entry (Mixed Effects Model – main effect of region: F(1,15200)=5.2, p=0.002; main effect of decision: F(1,15200)=101.1, p=8.5×10^-52^; region x decision interaction: F(1,15200)=11.6, p=5.3×10^-5^) and WZ exit (Mixed Effects Model – main effect of region: F(1,16655)=46.8, p=6.7×10^-13^; main effect of decision: F(1, 16655)=98.9, p=2.5×10^-24^; region×decision interaction: F(1, 16655)=63.6, p=1.3×10^-16^) LO ensembles burst upon entry in the WZ (Figure 5a). This burst was temporally aligned to the first tone presentation indicating a decrease in time remaining rather than the first tone presented in the WZ. Given that LO was active during OZ approach decisions and represented time remaining in the WZ, we hypothesized that the LO WZ entry burst would be disrupted for encounters that were quit. Interestingly, we found the opposite result — LO activity at WZ entry was higher during quits (Figure 5A; Mann-Whitney U Test - z=-3.31, p=9.4×10^-4^). This seemingly paradoxical finding suggests that anomalously high LO activity upon entering the WZ (and therefore exiting the OZ) might produce erroneous accept decisions even when the offer was bad. Consistent with this hypothesis, LO activity for offers with longer delays were significantly higher just prior to leaving the OZ when the offer would be quit compared to when it was either earned or properly skipped (S4A-B; ANOVA – F(2,2765)=7.7, p=4.6×10^-4^; post-hoc comparisons revealed more activity during quits than skips, p=0.002, and earns, p=0.002). Thus, heightened LO activity during quits further reflected the critical role that LO played in accept decisions. Although LO activity was elevated upon entering the WZ during quits, by the time rats quit the offer, LO activity was suppressed compared to leaving the WZ when earning reward (Figure 5B).

Aligned to WZ exit time, VO activity was elevated in the period leading up to a quit decision (Figure 5D). Elevated VO activity moments before leaving the WZ via a quit exceeded its reward-anticipatory activity when leaving the WZ via earning reward (Figure 5D; Mann-Whitney U Test = prior to WZ exit: z=-5.6, p=2.0×10^-8^). To determine when this elevated activity emerged, we compared VO activity between stays and quits oriented to WZ entry. Remarkably, VO discriminated stay from quit decisions immediately upon entering the WZ (Figure 5C). VO activity was elevated during late periods of the OZ that persisted into the WZ if the offer would be quit. In contrast, if reward would be earned, VO activity dipped below baseline levels upon WZ entry (Figure 5C; Mann-Whitney U Test = prior to WZ exit: z=-3.6, p=2.5×10^-4^). Decision encoding in VO remained when controlling for reward delays (S4D-E).

The temporal dynamics in LO and VO linked to WZ decisions suggested that LO activity dominated when reward would be earned whereas VO activity dominated when the offer would be quit. To further explore these opposing dynamics during WZ decisions, we measured the difference between LO and VO ensemble activity at each moment in time separately for stay and quit decisions. When rats earned reward, LO activity was elevated relative to VO activity upon entering and exiting the WZ (Figure 5E and 5F; Mann-Whitney U Test - WZ entry: z=7.09, p=1.2×10^-12^; WZ exit: z=5.6, p=1.4×10^-8^). In contrast, if the offer would be quit, VO activity was elevated relative to LO activity, and this difference became strongest in the moments leading up to leaving the WZ via quitting (Figure 5E and 5F; Mann-Whitney U Test – WZ exit: z=-2.03, p=0.04). Collectively, these results suggest a balance between LO and VO activity that guides WZ decisions, qualitatively similar to that observed in the OZ. LO participated in approach decisions that lead rats to rewards while VO participated in an override process that allowed rats to quit offers that were erroneously accepted. Finally, opposing representations in the WZ in LO and VO were uncorrelated on a moment-by-moment basis (Figure S4 F-G). These results further suggest that representational content in LO and VO are unlikely to directly implement a decision but are instead providing information to other structures that might do so.

As in the OZ, WZ decisions were driven by distinct valuation processes during the first 5 min of the session. Comparing WZ decision dynamics in LO and VO between early and later periods of the session can shed light on how these dynamics scaled with changes in the subjective valuation of offers as session time progressed. To address this, we compared ensemble activity during successive 5-min chunks of session time at WZ entry and exit for both stay and quit decisions. Unlike the burst of LO activity at OZ entry, the LO burst following WZ entry decreased as session time elapsed (Figure 5G and 5I; Mixed Effects Model – main effect of chunk at zone entry: F(10,1823)=7.04, p-6.1×10^-11^). This was true during both stays (Figure 5G-H) and quits (Figure 5I-J; Mixed Effects Model – no decision×chunk interaction: F(10,181)=1.13, p=0.33).

The first 5-min of session time was characterized by increased tendency to enter the WZ for all offers but subsequently quit those with a long delay. We predicted that LO activity would be elevated early in the session. This effect would follow from the notion that LO activity scaled with the subjective value of offers to guide approach decisions. Consistent with this possibility, LO activity was higher during the first 5-min of the session compared to the rest (Figure 5G-J; Mixed Effects Model – main effect of chunk: F(10,1830)=1.67, p=0.002). This was true during both stays (Figure 5G-H) and quits (Figure 5I-J; Mixed Effects Model - no decision×state interaction: F(10,1830)=1.67, p=0.08).

Given the role of VO activity in overriding approach decisions during quits, we further hypothesized that VO activity would be elevated during the first 5 min of the session. Consistent with this hypothesis, VO activity was elevated in the first 5 min of the session compared to the rest (Figure 5K-N; Mixed Effects Model – F(10,1370)=3.75, p=5.38×10^-5^). Increased VO activity early in the session occurred during both stays (Figure 5K-L) and quits (Figure 5M-N; Mixed Effects Model – no decision×chunk interaction: F(10,1360)=1.11, p=0.35). Thus, early periods of each session were characterized by a generalized increase in both LO and VO activity, reflecting heightened subjective value of both current and future rewards.

Although activity within each subregion changed across the session, the relative balance between them during both earns and quits was consistent across session time (Figure 5O-R). This result is consistent with that found in the OZ and suggests that LO and VO activity scaled with subjective value but their relative balance was conserved in the WZ.

### LO and VO represented specific features of encountered rewards

A characteristic neural signature of LO is heightened activity at reward delivery^67–70^, though the prevalence of reward-related activity in VO is unclear. To explore reward-related activity between subregions, we analyzed neural activity in three epochs surrounding reward: reward-anticipation (3-0 s prior to reward), reward-delivery (0-1 s following reward delivery), and reward-consumption (1-3 s following reward delivery). Reward-related activity differed between regions (Mixed Effects Model – main effect of region: F(1,635)=75.1, p=3.3×10^-18^) and epochs (Mixed Effects Model – main effect of epoch: F(1,635)=6.0, p=0.001; no region×epoch interaction: F(1,635)=0.14; 0.06). We therefore assessed neural activity in LO and VO separately between each reward epoch.

Prior to reward delivery, LO exhibited a rise in normalized firing rate, which peaked when sensory cues (sound of feeder) signaling imminent reward were presented (Figure 6A). VO exhibited a similar but weaker pattern as time to reward approached (Figure 6A; Mixed Effects Model – main effect of region: F(1,1913)=41.6, p=1.2×10^-11^; time×region interaction: F(1,1913)=10.3,p=0.0001). There was also a distinct activity pattern between LO and VO during reward consumption. VO ensembles dipped below baseline during consumption whereas LO ensembles remained elevated (Figure 6A). Thus, while reward anticipation and delivery were characterized by heightened activity in LO and VO, reward consumption was characterized by depressed activity only in VO.

**Figure 6.**
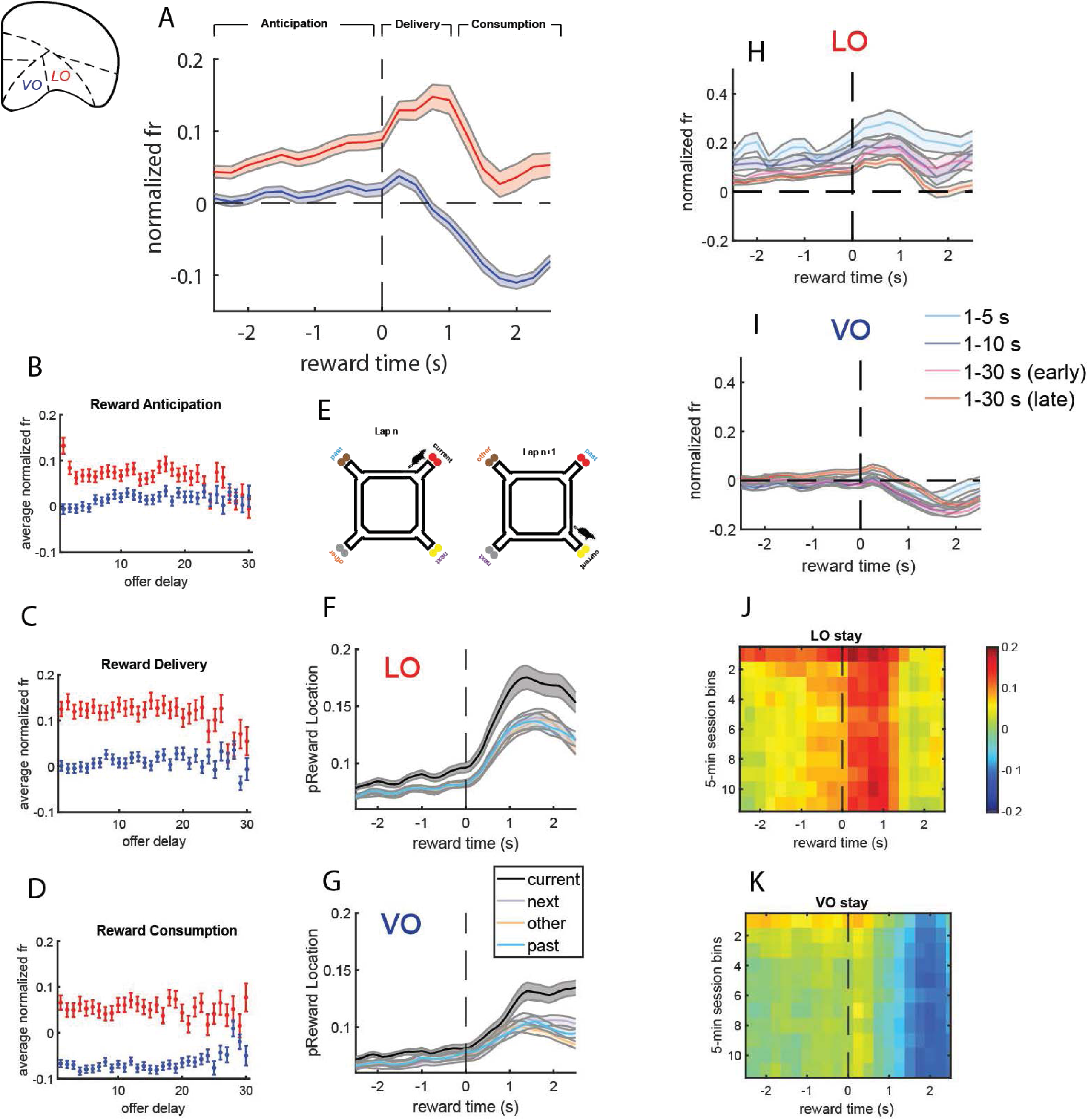
Reward-related activity in LO and VO adapted to economic and motivational factors. A) Mean (±SEM) neural activity within LO (red) and VO (blue) aligned to reward delivery. (B-D) Mean (±SEM) neural activity as a function of offer delay during anticipation (B), delivery (C), and consumption (D) in LO and VO. (E) Illustration of decoding analysis, current restaurant = location of the rat. Next, past, and other restaurants rotate with the rat’s current location to explore identity-specific representations. (F-G) Mean (±SEM) probability of decoding to each of the four rewards in LO (F), VO (G) during the epoch surrounding reward delivery. (H-I) Mean (±SEM) neural activity in LO (H) and VO (I) during reward across the four stages of training. (J-K) Mean neural activity in LO (J) and VO (K) broken up across successive 5-min session chunks.

Previous research has found that reward-related activity in the OFC is modulated by the delay preceding its delivery^71^. Inconsistent with previous work, reward delay did not influence neural activity in either LO or VO in any of the reward epochs (Figure 6 B-D; Mixed Effects Model – no delay×epoch interaction: F(2,16984)=0.16, p=0.08). Thus, retrospective information about offer delay when rewards were delivered was not encoded in LO or VO activity in Restaurant Row.

Reward-related activity within OFC represents specific reward features, such as flavor^72–78^. To explore the degree to which LO and VO represented specific reward features, we used a Bayesian algorithm to decode representations of reward flavor by restaurant and then reoriented the data relative to the restaurant the rat was currently at (Figure 6E). LO and VO ensembles encoded reward-specific information during anticipation (S5A-F; Wilcoxon Signed-

Rank Test – LO: z=10.66, p=1.6×10^-26^. VO: z=9.32, p=1.1×10^-20^; Mann-Whitney U Test on real vs. shuffled distributions – LO: z=5.26, p=1.4×10^-7^ VO: z=4.9, p=1.1×10^-5^), delivery (Wilcoxon Signed-Rank Test – LO: z=11.12, p=9.4×10^-29^; VO: z=8.77, p=1.7×10^-18^; Mann-Whitney U Test on real vs. shuffled distributions – LO: z=8.60, p=8.1×10^-18^; VO: z=5.27, p=1.4×10^-7^), and consumption (Wilcoxon Signed-Rank Test – LO: z=13.42, p=4.8×10^-41^; VO: z=14.31, p=1.9×10^-46^; Mann-Whitney U Test on real vs. shuffled distributions - LO; z=11.30, p=1.5×10^-29^; VO; z=14.22). Encoding of reward-specific information was significant across subregions prior to reward delivery, immediately after reward delivery, and during reward consumption (Figure 6F-G).

Given that decision and delay related activity adapted to economic conditions and motivational factors, we hypothesized that reward-related representations would also differ between training stages (global) and session time (local). To this end, we first compared neural dynamics surrounding reward delivery across the four stages of training. LO activity was reduced following introduction to the reward-scarce environment across all reward epochs (Figure 6H; Mixed Effects Model – main effect of stage: F(3,4356)=3.35, p=0.02; main effect of epoch: F(2,4356)=3.0, p=0.05; no stage×epoch interaction: F(6,4356)=0.79, p=0.54). Reduced reward-related LO activity was not linked to behavioral adaptations but instead emerged immediately when reward became scarce (Mixed Effects Model – post-hoc comparisons revealed a main effect on fixed coefficients during stage 3: t(4356)=-2.16, p=0.03; and stage 4: t(4356)=-2.77, p=0.005). In contrast, VO activity was unaffected by reward scarcity (Figure 6I; Mixed Effects Model – main effect of stage: F(3,6837)=1.63, p=0.18; main effect of epoch: F(2, 6837)=0.62, p=0.53; no stage×epoch interaction: F(6, 6837)=1.64, p=0.13), suggesting that reward-related modulations to economic conditions is a feature of more lateral aspects of the OFC.

Reward-related activity was also sensitive to motivational factors in both LO and VO. LO activity was elevated in the first 5 min of the session during reward anticipation (Mixed Effects Model – main effect of chunk: F(10,1390)=1.93, p=0.03) but not reward delivery (Mixed Effects Model – main effect of chunk: F(10,1390)=0.97, p=0.46) nor consumption (Mixed Effects Model – main effect of chunk: F(10,1390)=0.21, p=0.99). In contrast, VO activity was elevated early in the session across all three reward epochs (Mixed Effects Model – main effect of chunk during anticipation: F(10,1040)=5.33, p=1.0×10^-7^; delivery: F(10,1040)=2.12, p=0.02; consumption: F(10,1040)=3.26, p=0.0003). Thus, whereas all aspects of reward-related VO activity scaled with changes in subjective value across session time, only the reward anticipatory component of LO reward-related activity scaled with subjective value.

### Neural dynamics in LO and VO reflected distinct processes during initial valuations, reevaluations, and reward

Representational content and neural dynamics within LO and VO during OZ evaluations, WZ reevaluations, and reward delivery exhibited similarities and differences: LO exhibited bursts when cues signaling state transitions were presented and increased as reward became temporally proximal. VO showed elevated activity during long delays and was elevated in the moments leading up to decisions that resulted in rats abandoning current rewards via skips or quits. These qualitatively similar dynamics within each subregion might reflect the same process at distinct task stages. However, previous data suggests that distinct decision-making processes drive decisions in the OZ and WZ^49,50,53–56.^ To test whether neural dynamics within OFC subregions reflected similar or dissimilar processes in the WZ and OZ, we examined the degree of overlap in the cells participating in key neural signatures within LO and VO throughout task states. For LO, we assessed the overlap in cells participating in bursts aligned to OZ entry, WZ entry, and reward delivery. For VO, we assessed the overlap in cells participating in leaving a task zone during skips, quits, and leaving the restaurant after earning reward (i.e., “linger zone leaving”).

There was little overlap in LO cells and little correlation in firing rates during OZ and WZ entry bursts (Figure 7D; OZ-sensitive cells: Pearson’s Correlation: r=.13, p=.063; WZ-sensitive cells: Pearson’s Correlation: r=-.11, p=.92). Interestingly, there was a negative correlation in activity scores at OZ entry and reward delivery (Figure 7E; r=-5.9, p=1.9×10^-14^), suggesting that OZ cells tended to be depressed at reward delivery. We did not find a significant correlation in activity at OZ entry and reward delivery for cells sensitive to reward delivery (Figure 7E; Pearson’s Correlation: r=-.04, p=.61). Finally, there was no significant correlation in activity at WZ entry and reward delivery for either WZ-sensitive cells (Pearsons Correlation: r=.05, p=.27) or reward-delivery-sensitive cells (Figure 7F; Pearsons Correlation: r=.006, p=.46). This negative correlation was also evident in the peri-event averages, which revealed that cells that were sensitive to a given zone entry were only weakly modulated (if at all) at the other two zone entries (Figure 7A-C). Thus, distinct clusters of LO cells marked transitions into the OZ, WZ, and reward delivery. This result is consistent with prior observations of distinct LO cells marking state transitions in appetitive learning tasks^67,68^.

**Figure 7.**
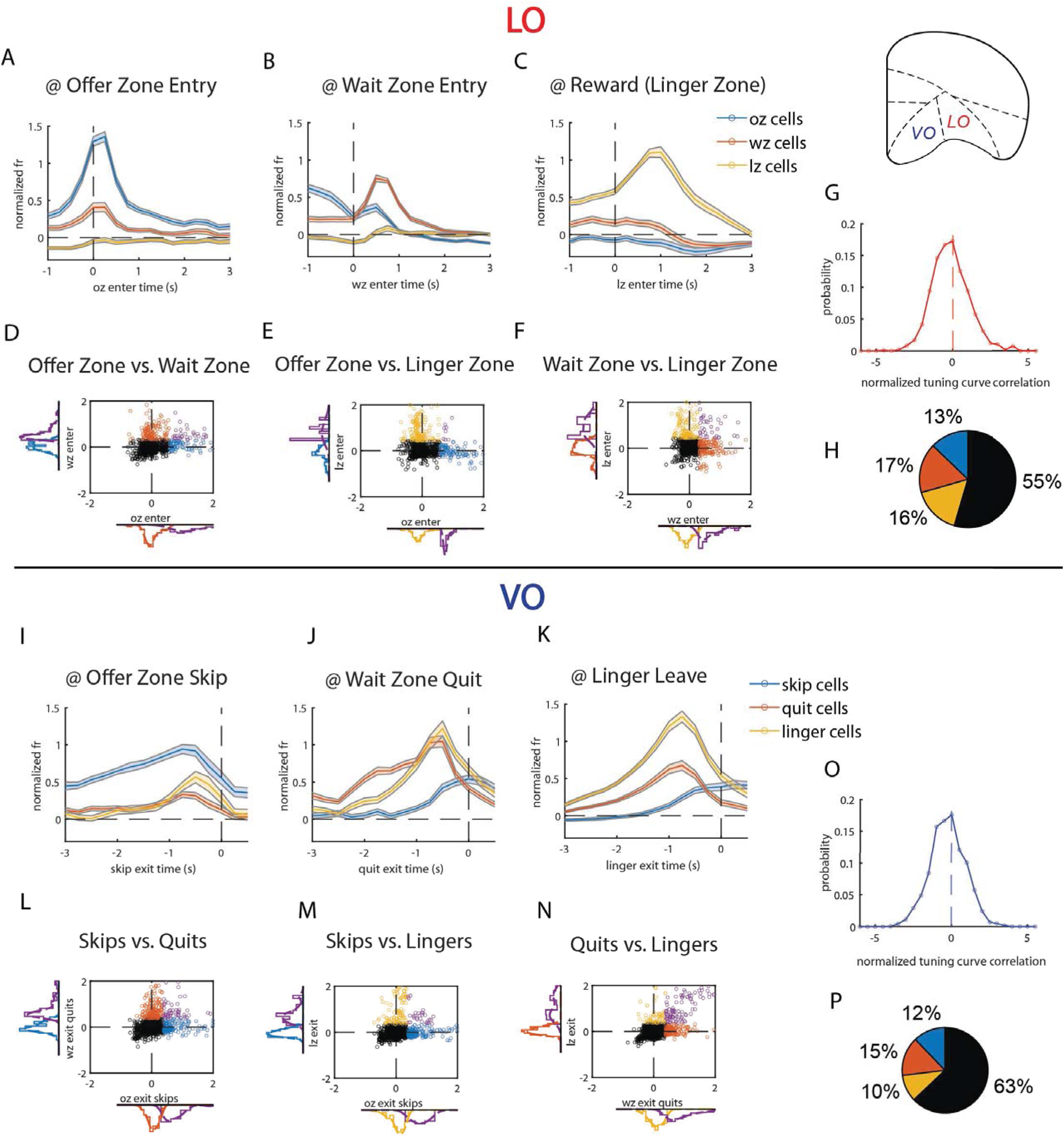
Distinct clusters of cells participate OZ evaluations, WZ re-evaluations, and reward. (A-C) Mean (±SEM) normalized firing rate plotted at OZ entry (A), WZ entry (B), and reward delivery (C) for LO cells sensitive to OZ entry (blue), WZ entry (orange), and reward delivery (yellow). (D-F) Scatter plots of normalized activity scores for LO cells. (D) Mean normalized activity of LO cells at OZ (x-axis) and WZ (y-axis) entry, (E) at OZ entry (x-axis) and reward delivery (y-axis), (F) and at WZ entry (x-axis) and reward delivery (y-axis). Black = not sensitive to either zone entry event; blue = sensitive to OZ entry; orange = sensitive to WZ entry; yellow = sensitive to reward; purple = sensitive to events that define both axes. (G) Probability distribution of within-cell correlations of tuning curves to delay in the offer and wait zone within the LO. Correlations were normalized relative to a shuffled distribution in which the relation between normalized firing rate and delay were randomized. (H-N) Equivalent layout to (A-F), but for VO cells sensitive to skipping in the offer zone (blue), quitting in the wait zone (orange), or leaving in the linger zone (yellow).

While these results suggest LO bursts and zone entry were distinct, it remains possible that LO cells encoding information about reward delay did so similarly between the OZ and WZ. By comparing shuffle-normalized correlations in tuning curves to delay in the OZ and time-remaining in the WZ for each cell, we found no significant relation between delay-sensitive activity between the two zones (Wilcoxon Signed Rank Test: real versus shuffled distributions, z=1.2, p=.25; Figure 7G). Collectively, these results suggest that LO ensembles participated in distinct computational processes throughout task states in Restaurant Row, rather than reflecting a singular process that was reactivated during sequential decisions in the OZ and WZ.

VO activity was heightened not only in the moments leading up to skips and quits, but also in the moments leading up to leaving the linger zone (Figure 7J). We therefore assessed the degree of overlap in the collection of cells that participated in leaving a restaurant via a skip, quit, or lingering. There was no significant correlation in average activity prior to skipping and quitting for cells sensitive to either skips (Figure 7K; Pearsons Correlation: z=-.004, p=.52) or quits (Figure 7K; Pearsons Correlation: z=.04, p=.30). In addition, peri-event averages of normalized activity revealed that quit-sensitive cells exhibited less pronounced activity prior to skipping than skip-sensitive cells (Figure 7H). Conversely, skip-sensitive cells exhibited less pronounced activity prior to quitting than quit-sensitive cells. Finally, we did not find a significant correlation between tuning curves within VO cells to reward delay in the OZ and WZ (Figure 7n; Wilcoxon Signed Rank Test: real versus shuffled distributions: z=-.73, p=.46). Thus, the decision-related processes that VO participated in during OZ skips were distinct from the processes that implemented WZ quits.

Surprisingly, there was substantial overlap in the cells that fired prior to quitting out of the WZ and leaving after lingering (Figure 7I, 7J, and 7M). Average activity of quit-sensitive cells when leaving a restaurant prior to quitting and lingering were significantly correlated (Figure 7M; Pearsons Correlation: r=.47, p=2.9×10^-14^). Average activity of linger-sensitive cells when leaving a restaurant during lingering and quitting were significantly correlated (Figure 7M; Pearsons Correlation: r=.61, p=6.4×10^-18^). When comparing neural dynamics of each cell type prior to quitting, activity in cells that were sensitive to leaving a restaurant following post-reward lingering ramped up and peaked in a manner similar to quit-sensitive cells (Figure 7I). Conversely, activity in quit-sensitive cells ramped in the moments leading up to a linger-leave decision, though the degree and peak up ramping was weaker than it was in linger-leave-only cells (Figure 7J). Thus, compared to the distinct collections of VO cells that participated in skips and quits, there was substantially more overlap in the cells that participated in quits and leaving after reward.

Overall, these results are consistent with distinct decision-making mechanisms deployed in the OZ and WZ, with opposing processes implemented in LO and VO. While LO ensembles participated in approach-related behaviors in both zones (i.e., accepts and stays), they did so as part of distinct decision-making processes. VO ensembles participated in leave-related behaviors in both zones (i.e., skips and quits), but they did so as part of distinct decision-making processes. However, there was overlap in the processes participating in quitting an offer (prior to reward) and leaving a restaurant following a lingering period (after reward). Thus, VO ensembles appeared to participate in one process when leaving a restaurant while being spatially distant from reward (OZ skips) and a separate process when leaving a restaurant while being closer to reward (quits and linger-leaves).

## Discussion

Lateral and ventral aspects of the OFC exhibited marked differences in representational content and timing during sequential foraging decisions. LO encoded information about the subjective value of a current reward whereas VO encoded information about the opportunity costs associated with pursuing it. These representations tracked approach/leave decisions and adapted to external, economic factors along with internal, motivational factors. Thus, lateral and ventral aspects of the OFC contained opposing representations to guide sequential foraging decisions.

Despite distinct representations in LO and VO, two important observations from the present study suggest a shared coding scheme between subregions in which information about the subjective value of reward and the underlying state of the task (i.e., OZ, WZ, and LZ) were multiplexed. First, opposing representations were preserved along the sequence of events that defined an encounter (i.e., “task states”), but with distinct clusters of cells implementing them within each subregion across task states. This result is consistent with the observation that OFC ensembles encode goal locations throughout a spatiotemporal trajectory to reward^72^. Second, overall activity in both subregions scaled with changes in subjective value induced by external changes in reward availability and internal changes in reward motivation. Collectively, these two observations are consistent with the notion that OFC ensembles multiplex information about subjective value and task state^72,74,79^. Information related to subjective value (i.e., current reward in LO and opportunity costs in VO) was encoded via overall ensemble activity whereas information about location along a reward trajectory (i.e., task state) was encoded via distinct clusters of cells. This multiplexed coding scheme integrates two influential theories of OFC function, one of which suggests that OFC represents the subjective value of rewards^10,58,60,61^ and that the other that OFC represents task state^13,79^. Results from the present study, combined with other empirical and theoretical work^74,79^, suggest that these are not mutually exclusive possibilities but are instead realized as distinct features of neural activity in the OFC.Importantly, however, results from the present study illustrate that while this coding scheme was shared between lateral and ventral OFC, the precise content and timing of representations to which they apply differed between subregions.

The OFC integrates information across environmental dimensions (e.g., sensory cues, spatial locations, memories) to guide behavior^31,57,58,60,61,71,72,80^. Results from the present study suggest a novel mechanism by which information within distinct OFC subregions might be integrated to guide behavior. Rather than integrating information across environmental dimensions within a single subregion of the OFC, the present study revealed distinct information in distinct OFC subregions. Cells in lateral OFC (LO) provided information needed to make approach decisions while cells in ventral OFC (VO) provided information needed to forgo a reward and continue searching. Because the balance of LO and VO signals tracked decisions, relative activity in LO and VO between approach/leave decisions provide an “integrated signal” for sequential foraging decisions.

While the integrative signal in relative LO-VO activity was correlated with approach/leave decisions at each stage of an encounter, it is unlikely to reflect an evidence accumulation signal that implements the decision itself. This possibility is precluded by the observation that LO and VO ensemble activity was uncorrelated on a moment-by-moment basis. Thus, it is more likely that LO and VO provide independent sources of information that are used by downstream structures to transform value-related representations into decisions. One candidate structure to this end would be the anterior cingulate cortex (ACC). Previous work exploring information processing across the medial wall in rodents using Restaurant Row found that ACC exhibits gradual ramping in the moments leading up to a choice^49^. In addition, research using patch-foraging tasks has shown that ACC activity ramps until reaching a threshold to trigger a decision to leave a currently exploited patch and begin searching elsewhere^81^. Thus, OFC ensembles might provide value-related and task-state related information that are sent to ACC which accumulates evidence in favor of a decision. This is precisely what Belewski et al., (2023) found with simultaneous OFC/ACC recordings in primates making value-based decisions. Our results further raise the possibility that distinct subregions within the OFC provide opposing representations that are needed to move the ACC needle in one direction or another.

Results from the present study suggest that foraging decisions are implemented by multiple decision-making mechanisms in different stages of the task^82^. Previous work in Restaurant Row^49,50,53–56^, along with work using other sequential foraging tasks^83^, similarly suggests that multiple decision-making mechanisms are deployed when animals forage for sequentially-encountered rewards. Importantly, we observed distinct decision-making mechanisms during sequential foraging in Restaurant Row were realized in distinct OFC subregions. Initial evaluations were governed by opposing processes that supported OZ decisions. The fast-acting accept process was reflected in heightened LO activity that encoded subjective benefits associated with the current reward. The skip process was slower, overrode prepotent approach tendencies, and was reflected in ramping activity that scaled with opportunity costs in VO. Interestingly, formal models of foraging suggest precisely these representations to make optimal decisions - the profitability of the current offer should be compared to the opportunity costs associated with pursuing that offer to make an accept/reject decision^63–66,84^. Thus, results from the present study suggest that these opposing representations are realized in LO and VO as part of conflicting processes that make evaluations to either accept a reward or continue searching.

The operation of these dual processes also appeared to give rise to peculiar suboptimalities in how rats made decisions in Restaurant Row due to the unintuitive temporal dynamics of how opportunity costs were represented in the VO. The most time-efficient strategy would be to go directly to the WZ to initiate the countdown to reward and then decide to stay or leave (e.g., rather than spending time outside of a restaurant deciding whether to enter and wait in line, it is best to just get in line and then decide if it is worth continuing to wait). Participants across species do not exhibit this decision-making pattern in Restaurant Row^49,53,55,85–87^. In addition, because opportunity costs are known prior to each encounter, there is no need for a neural representation of opportunity costs to gradually ramp as if it were accumulating evidence. Nevertheless, a suboptimal, ramping representation of opportunity costs was observed in VO ensembles. Perhaps the ramping nature of opportunity cost representations in VO reflects an increasing urgency to make a decision, similar to collapsing boundaries in drift diffusion models of decision making.

Evidence from the present study on heterogeneity between LO and VO is correlational. Future research is needed to clarify functional similarities and differences between these OFC subregions. Nevertheless, results from the present study lend predictions for how disruptions to lateral vs. ventral aspects of the OFC should affect behavior in sequential foraging tasks.

Specifically, disrupting VO ensembles should impair the ability to reject currently offered rewards and continue searching for future prospects. This is consistent with existing evidence showing that ventral aspects of the OFC are involved in behaviors that are temporally distant from reward and require extended motivation to achieve goals. Based on results from the present study, a similar result should be found by artificially increasing activity in lateral aspects of the OFC. Heightened activity in the LO would enhance representations of the current offer’s value and therefore result in increased propensity to approach currently offered rewards. Thus, results from the present study suggest that disrupting the relative balance of LO-to-VO activity should have specific effects on sequential foraging decisions.

Results from the present study are unlikely to reflect purely motoric signals that encode movement parameters. First, OFC subregions exhibited drastically different levels of activity in the WZ based on 1) which decision would be made, and 2) before and after reward was delivered, even though rats were in the same spatial location that imposed the same movement constraints. Similarly, delay-dependent activity in the OZ was more consistent with a cognitive interpretation rather than a motoric one given the fixed and relatively constrained spatial location and movement opportunities in the OZ. Third, activity in each subregion was affected by contextual variables determined by both external factors (i.e., reward scarcity) and internal factors (i.e., hunger) despite being confined to the same maze that imposed the same constraints on movements.

Unlike the present study, previous research directly addressing OFC heterogeneity in rodents has focused on distinctions between medial and lateral aspects of the OFC. Malvaez et al., (2020) found that ventrolateral aspects of the OFC were critical for learning and encoding reward value whereas medial OFC was critical for retrieving reward value to inform goal-directed decisions. Lopatina et al., (2017) found that lateral OFC encoded information about the sequence of events leading up to reward whereas medial OFC encoded information about reward value that generalized across the sequence of events leading to reward. Finally, Hervig et al., (2020) found that disrupting lateral OFC decreased sensitivity to feedback and therefore increased perseverative behavior during reversal learning. Disrupting medial OFC increased sensitivity to feedback and therefore improved performance during reversal learning through decreased perseveration.

Our finding that LO activity was heightened during approach to reward via distinct ensembles in an experience-dependent manner is consistent with this previous research showing that LO is critical for learning about reward value^20^, marking transitions to reward^19^, and is required for reward-based feedback^22^. Results from the present experiment expand these findings by suggesting that LO plays a role in learning, sensitivity to feedback, and marking key transitions in reward approach by encoding information about the subjective value of a prospective reward during approach decisions. However, our finding that VO activity represented opportunity costs through ramping activity that guided decisions to leave a prospective reward is inconsistent with previous research showing that medial aspects of OFC are required to retrieve learned reward value during approach decisions and is critical to reward-based feedback. This inconsistency stresses the important point that VO ensembles in rodents participates in distinct processes from both more lateral and medial aspects of the OFC and should therefore be treated as a distinct subregion within the OFC.

## Acknowledgements

We would like to thank Chris Boldt and Ayaka Sheehan for their invaluable contributions to surgery and histology, respectively. In addition, we would like to thank Geoff Diehl and the rest of the members of the Redish lab for helpful feedback throughout the project. This work was supported by NIH training grant T32DA037183.

## Author Contributions

P.J.C and A.D.R designed the study while P.J.C (along with Chris Boldt and Ayaka Sheehan) collected data and supported technical aspects of the project. P.J.C and A.D.R conceived and implemented neural and behavioral analyses. Finally, P.J.C and A.D.R wrote the manuscript.

## Methods

### Experimental Model and Study Paricipant Details

#### Subjects

Eight adult Fisher-Brown Norway rats served as subjects (4F, 4M). Because we did not obtain large cell yields from one of the rats (R665), his data was excluded from all analyses.

Thus, the effective sample for the analyses presented in this paper was n = 7 (4F, 3M). Rats were single-housed in a temperature-controlled colony room with a 12 hr light/dark cycle. Each rat’s daily food intake was obtained from the food earned within hour-long experimental sessions.

Rats had free access to water in their home cages. All procedures were approved by the University of Minnesota Institutional Animal Care and Use Committee (IACUC).

#### Histology

Following completion of recording sessions, rats were euthanized via pentobarbital overdose and were then perfused. Probes were removed from the brain following perfusion. Brains were then removed and post-fixed in paraformaldehyde for 24 hours, after which they were placed in a jar containing a mixture of paraformaldehyde and sucrose for cryoprotection.

Coronal sections (40 microns) were cut using a cryostat and were subsequently stained with cresyl violet. Probe trajectory across coronal slices, coupled with records of surgical placement and probe turning were used to confirm probe placements within the OFC.

#### Surgery

Rats were anesthetized with 2% isoflurane using an induction chamber prior to being placed on the stereotax. A custom-built head ring was attached to the skull using Metabond.

The dura was removed and three craniotomies were drilled through the skull. One craniotomy was drilled directly above the ventral orbital area (VO; A/P = 4.2, M/L = 1.5) while another was drilled directly above the lateral orbital area (LO; A/P = 4.2, M/L = 2.5). Craniotomy placements were determined using the Paxinos and Watson atlas. Hemisphere placement for each OFC subregion was counterbalanced across subjects. For each rat except R581, three 32-channel Cambridge Neurotech E-2 silicon probes, each containing two shanks (9mm long with 250 microns of spacing between them), were then lowered into the cortex, one probe per craniotomy. For R581, two 64-channel Cambridge Neurotech F silicon probes were used, with 6 shanks per probe (6mm long with 200 micron spacing between each). Probes were connected to custom-built hyperdrives to allow for turning and localization. Each probe was lowered 1-1.5 mm ventral during surgery. Once probes were in place, craniotomies were filled with bone wax.

The microdrives were then attached to the skull using Metabond. Finally, a custom-made skull ring and headcap were attached to the head ring to protect the implant. Following surgery, probes were turned down in small increments each day until reaching their terminal location. Most turning took place prior to recording sessions and during the first 10 recording sessions.

#### Behavior Training

Prior to behavioral sessions on Restaurant Row (RRow), rats were given 30 min of access to one of the four flavored pellets (plain, chocolate, cherry, banana; 45 mg per pellet) each day until they readily consumed each flavor. Once rats were familiarized with each flavored pellet, they were trained to run clockwise around the experimental track to earn reward (2 flavored food pellets per encounter) at each restaurant during a 60 min session, conducted once per day. During the early phase of pre-training, rewards were delivered 1 s after entering the restaurant, and this delay range increased across sessions. Specifically, once rats reliably ran the maze to earn rewards, the delay range increased to 1-5 s. Following 7 sessions of training with 1-5 s, the delay range increased to 1-10 s, which was in effect for another 7 sessions. The purpose of this training regime was to ensure that rats approached each restaurant, in turn, and earned enough rewards to fulfill their daily food requirements. Note that pre-training stopped at a delay range of 1-10 s.

#### Behavioral Recording Sessions

To ensure rats were comfortable running on the maze while plugged in to the recording equipment, rats ran up to five sessions of RRow with a 1 s delay to reward. Following this brief retraining period, experimental recording sessions began. As in training, each session lasted 60 min. In addition, rats were placed on a flowerpot for a 5 min period before and after experimental sessions to collect neural data during rest. There was a total of 35 recording sessions for each rat except R621 (23 sessions) and R626 (24 sessions), both of which ended early due to infections. The first 5 recording sessions consisted of the 1-5 s delay range (stage 1). The second 5 sessions (i.e., sessions 6-10) consisted of the 1-10 s delay range (stage 2). The final 25 sessions consisted of the 1-30 s delay range (stages 3-4). The final 25 sessions of the task is referred to as the “reward scarce environment” in which rats must decide how to allocate their limited time to earn their needed daily food intake. Importantly, this was the first time rats experienced such long delays in RRow.

### Method Details

#### Neural Data Acquisition and Processing

Neural data were collected using a 256 channel Intan RHD 2000 system. Each rat was plugged into the Intan system via two headstages; a 64 channel headstage that was connected to the two-probe microdrive and a 32 channel headstage plugged into the single-probe microdrive. Voltage signals at each channel were sampled at 30 kHz.

Kilosort 2.0 was used to identify single-unit clusters within each recording session (including time on the flowerpot before and after the behavioral session). Voltage signals from each probe were normalized via median subtraction to eliminate noise and went through a 600 Hz high-pass filter to isolate potential spikes. Finally, sorting results for each probe and each session were manually curated using Phy. Each probe was analyzed separately to identify clusters.

### Quanificaion and Staisical Analysis

The primary statistical technique used for analyses was a mixed effects ANOVA. This method was used because it provides a flexible and robust way to explore multidimensional main effects, along with interactions of hierarchically nested data and potentially correlated predictor variables^10,7^. All mixed effect ANOVAs for neural analyses were performed on data averaged across cells within a single session and then concatenated across sessions. The specific formula used for each analysis is presented with the text corresponding to that analysis (below). One-Way ANOVAs were used to assess simple main effects for behavior. Finally, Mann-Whitney U tests and Wilcoxon rank sum tests were used to assess whether distributions of behavioral or neural quantities were significantly different from each other or whether they were significantly different from a zero-median distribution, respectively. Bonferroni corrections for statistical significance tests were used to adjust for multiple comparisons. Specification of each statistical methods application to a given analysis is specified below.

*Defining economic conditions (training stages):* Recording sessions were divided into four stages to explore how behavioral and neural measures adapted to a reward scarce environment. The first stage consisted of the first 5 sessions of recording with a 1-5 s delay range. The second stage consisted of the next 5 sessions with a 1-10 s delay range. The third stage was the first 7 sessions of the 1-30 s delay range while the fourth stage consisted of the final 18 recording sessions with a 1-30 s delay range. The third stage reflected early periods of the reward scarce environment in behavior was in progress of adapting the new conditions. The fourth stage reflected later periods of the reward scarce environment (last 18 days) in which behavior adapted and stabilized. These training stages reflect global economic conditions because they determine the long-term availability of food (on the scale of days) within a given training stage. Behavioral and neural adaptations to the global economic climate were assessed through comparisons between these four training stages.

*Defining motivational factors (chunked session time):* Rats exhibited distinct behaviors during the first 5 min of each session compared to the remaining 55 min. To measure these changes, we binned behavioral and neural data into successive 5-min chunks within a session. These 5- min chunks reflect the local economic climate because they are defined by transient and brief periods (on the scale of minutes) in which rats valued food differently. While the economics on a global scale are determined by reward availability as set by the experimental protocol (i.e., externally determined), economics on this more local scale are presumably determined by motivational factors (i.e., internally determined) and were transient and not an inherent aspect of RRow or the experimentally imposed training sequence. Behavioral and neural changes to the local economic climate were assessed through comparisons across successive 5-min chunks of session time.

*Figure 1 Behavior Analyses:* Because there were no significant differences in the subjective value across restaurants, all behavioral analyses were performed on encounters concatenated across restaurants within a session.

A lap was defined as an encounter with an active restaurant. The rate at which laps were completed across session time was measured as the number of laps during each minute of the session. Lap rates were concatenated across sessions and grouped into the four training stages to determine how within-session changes in lap rate were affected by reward scarcity. A Mixed Effects Model (2 fixed effects and 1 random effect) was used to quantify changes in lap across session time and stages: lap rate ∼ sessiontime*stage + (1|ratID).

Time spent making a decision in the OZ was measured as the difference between the time at which the offer zone was exited and the time at which the offer zone was entered for each encounter. Time spent in the OZ was analyzed separately between accepts and skips. A Mann-Whitney U-test on the distribution of time spent making accept and skip decisions across laps concatenated across all sessions was used to compare decision-dependent differences in OZ decision time. To explore how the global economic climate influenced decision time, OZ time spent was averaged across all laps for each session, and sessions were grouped into the four training stages. A repeated measures ANOVA with training stage as a factor was used to test the effect of the global economic climate on OZ decision time. OZ decision time was also analyzed across reward delays conditioned upon each decision. Time spent in each trial was normalized relative to the average time spent in the OZ for that session to normalize between-rat differences in overall, average time spent. A repeated measures ANOVA, with decisions and delays as factors, was used to test for effects of decisions and delays on time spent in the OZ.

Linger time was defined by the amount of time rats spent at the current restaurant after receiving reward and prior to leaving for the next restaurant. This metric was calculated for each lap by taking the time at which the restaurant was exited from the time at which reward was delivered. Linger time was averaged within each session, and sessions were then grouped into the four training stages to assess how reward scarcity affected linger time. A repeated measures ANOVA with training stage as a factor was used to test the effect of the global economic climate on linger time.

Sensitivity of OZ and WZ decisions were measured as the probability of skipping (OZ) or quitting (WZ) an offer as a function of delay within each session. Sessions were grouped into the four training stages to determine how the psychometric choice curve was affected by the global economic climate. A repeated measures ANOVA, with delay and task stage as factors, was used separately for OZ and WZ decisions to test for effects of offer delay and training stage on choice.

The number of quits and skips were binned into successive 5-min chunks within each session during stages 3 and 4 to determine how decision frequencies changed with the local economic climate. Decision counts within each bin were then averaged across sessions. A repeated measures ANOVA, with session chunk as a factor, was used to test for the effect of session time on decision frequency.

Time spent making decisions in the OZ for all encounters were binned into successive 5- min chunks within each session to determine how OZ decision time changed with the local economic climate. Only data from stages 3 and 4 were used for this analysis. Time spent in the OZ was that concatenated across sessions to measure the probability density function of time spent within each 5-min chunk. A repeated measures ANOVA, with lap chunk as a factor, was used to test for the effect of session time on OZ decision time.

To measure how sensitivity of OZ and WZ decisions to delay changed with local economic conditions, choice data were binned into successive 5-min chunks and a threshold indicating how long rats were willing to wait for reward was measured within each chunk. To this end, within each 5-min chunk for each session from stages 3 and 4, a Heaviside function was fit to the distribution of binary choices and corresponding delays to estimate the delay at which the probability of making a choice was 0.5. These thresholds were then averaged across sessions. A repeated measures ANOVA, with session chunk as a factor, was used separately for OZ and WZ decisions to test how sensitivity to delay changed across session time.

*Figures 2-3: Offer Zone Evaluations.* Each cell’s firing rate was normalized by subtracting its firing rate in each temporal bin (250 ms) by the cell’s session-wide mean firing rate and dividing by that cell’s session-wide standard deviation in firing rate. All event-related measures of neural activity used this z-scored firing rate. Finally, because there were no significant differences in he subjective value across restaurants, all neural analyses were performed on encounters concatenated across restaurants within a session. A window of -0.5 to 2.5 s around OZ entry time was used while a window of -2.5 to + 0.5 s was used around OZ exit time to assess neural dynamics at key behavioral moments. These temporal parameters were determined based on observations of typical occupancy time in the OZ across rats.

To compare neural dynamics between LO and VO in the OZ, normalizing firing rate was averaged across cells within each OFC subregion across all encounters within each session (resulting in a sessions × regions × time matrix). Only sessions from stages 3 and 4 were used. A mixed effects model (2 fixed effect and 1 random effect) was used to test for region- and time-dependent differences in OZ neural activity: fr ∼ time*region + (1|ratid).

To measure how neural activity changed as a function of reward delay as time in the OZ unfolded, normalized firing rate for each cell was averaged across encounters for each delay within the session. Normalized firing rate was then averaged across cells within each subregion, resulting in a sessions × regions × delays × time matrix. A mixed effects model (2 fixed effect and 1 random effect) was used to test how ensemble activity changed with offer delay and time: fr ∼ time*delay + (1|ratid).

Tuning curves of normalized firing rate to reward delay for each cell were constructed to further determine sensitivity to offer delay within each subregion. Because neural dynamics observed from event-related averages revealed time-dependent sensitivity to delay, tuning curves to reward delay were constructed separately during OZ entry and during the putative decision period. Tuning curves to delay at OZ entry were constructed using data only from the first 1 s of time spent in the offer zone (i.e., 0-1 s after offer zone entry). Tuning curves during the putative decision period were constructed from data excluding the first 1 s of time spent in the offer zone. Tuning curves were then averaged across cells within each session (only stages 3 and 4), resulting in two separate sessions × regions × delays matrices of normalized firing rates, one for each epoch in the OZ. A mixed effects model (1 fixed effect and 1 random effect) was used for both OZ entry and decision time tuning curves to test for an effect of delay in each subregion: fr ∼ delay + (1|ratID).

Ramping activity within time spent in the OZ was measured as the slope of normalized firing rate to time spent in the offer zone. To this end, normalized firing rate for each cell was averaged across all encounters of a given delay and within a given range of time spent in the OZ, resulting in a cells × delays × time-in-offer-zone matrix for each session (only stages 3 and 4). A line was then fit to normalized firing rate for each cell at each delay × time-in-offer-zone combination. Slopes were subsequently averaged across time spent in the OZ to yield a cells × delays matrix which quantified how steeply neural activity rose with time in the OZ across delays for each cell. Finally, the slopes across cell were averaged for each session. A Mixed Effects ANOVA (1 fixed effect and 1 random effect) was used to test the effect of delay on slopes (acceleration of neural activity) within each subregion: fr ∼ delay + (1|ratid).

To examine the development of offer zone ramping activity in VO across training stages, the slope relating neural activity to time spent in the OZ were averaged across delays within each session. The slopes relating neural activity to OZ time were then averaged across sessions within each training stage. A mixed effects model (1 fixed effect and 1 random effect) was used to test the effects of training stages on second-order slopes: fr ∼ stage + (1|ratID).

The emergence of delay-dependent ramping in VO across the reward scarce environment was measured by finding the slope relating ensemble activity within each session to offer delay. We then fit a line to obtain a second-order slope that measured the degree to which VO ramping activity (which was itself measures as the slope of activity across time spent) was sensitive to delay. A second-order slope of 0 would mean that VO ramping activity was unrelated to delay, whereas a slope > 0 means that VO activity accelerated more steeply for longer delays. We then averaged this second order slope across sessions within each training stage. A mixed effects model (1 fixed effect and 1 random effect) was used to test the effects of training stages on second-order slopes: fr ∼ stage + (1|ratID).

To examine decision-dependent neural dynamics in LO and VO as time in the OZ unfolded, average normalized firing rates for each cell were obtained separately during accept and skip encounters. Normalized firing rates were then averaged across the temporal windows surrounding OZ enter and exit times to yield a single value reflecting average neural activity between decisions for each cell. Finally, activity scores were averaged across cells within each session (only stages 3 and 4), resulting in a sessions × decisions matrix of average ensemble activity during OZ entry and exit. Mann-Whitney U tests on the distributions of average normalized firing rates were used to determine decision-dependent differences in neural activity between LO and VO.

To further clarify how relative activity in LO and VO changed throughout the OZ between decisions, we measured the difference in ensemble activity between LO and VO at each moment in time during accept and skip encounters within each session. Specifically, an encounters × time × decisions matrix of cell-averaged normalized firing rates for each session (only stages 3 and 4) was constructed. We then subtracted this matrix for LO ensembles from that for VO ensembles and averaged this matrix across encounters to obtain a sessions × time × decisions matrix of LO minus VO normalized firing rates. A Mann-Whitney U test on the distribution of LO-VO scores averaged across the temporal windows at OZ enter and exit times was used to test for decision-dependent differences in relative LO-to-VO activity.

To examine how decision-dependent activity changed to local economic conditions within session, neural activity for each cell was averaged across all quit and stay encounters during each successive 5-min bin for each session (only stages 3 and 4). Normalized firing rate was then averaged across cells within each 5-min bin, resulting in a sessions × chunks × decisions matrix. A mixed effects model (2 fixed terms and 1 random term) was used to test for the effects of decision and session time on neural activity: fr ∼ chunk*decision + (1|ratID).

*Figure 4-5: Wait Zone Representations and Re-evaluation.* To examine sensitivity to time remaining and time spent, 3-dimensional (time remaining × time spent × initial delay) tuning curves of normalized firing rates were constructed for each cell in each session (only stages 3 and 4). Tuning curves were then averaged across cells within each subregion for each session, resulting in a sessions × time remaining × time spent × initial delay matrix. A mixed effects model (3 fixed effects and 1 random effect) was used to test for significant encoding of WZ temporal intervals: fr ∼ timeSpent*timeRemaining*initialDelay + (1|ratID).

Decision-dependent neural dynamics within each subregion was analyzed using event-related averages of neural activity during quit and stay encounters at WZ entry and exit. To this end, normalized firing rate was averaged separately across stay and quit encounters for each cell and each session (only stages 3 and 4). Normalized activity was then averaged across cells within each subregion for each session, resulting in two sessions × time × decisions matrices, one for WZ entry and the other for exit. Finally, ensemble activity was averaged across the temporal windows during entry and exit time to yield a single value measuring average ensemble activity between WZ decisions. In addition, the difference between cell-averaged LO and VO at each moment in time was measured for each session (only stages 3 and 4) between stay and quit decisions. Relative LO-to-VO scores were also averaged across temporal windows at WZ entry and exit. Mann-Whitney U tests on the distributions of normalized firing rate and relative LO-to-VO activity were used to determine differences in neural activity between LO and VO during earns and quits.

To examine how decision-dependent activity changed to local economic conditions within session, normalized firing rate was averaged across stay and quit encounters in successive 5-min intervals for each cell at WZ entry and exit. Neural activity in each 5-min interval was then averaged across cells for each session (only stages 3 and 4). The difference between cell-averaged LO and VO activity at each moment in time during earns and quits was also measured for each session to explore how relative LO-to-VO activity changed across session. A mixed effects model (2 fixed terms and 1 random term) was used to test the effects of decision and session time on both normalized firing rate within each subregion: fr ∼ chunk*decision + (1|ratID), and relative LO-to-VO activity: LO-VO ∼ chunk*decision + (1|ratID)

To examine the decision-dependent burst of LO activity following WZ entry, cell-averaged normalized firing rate was measured during the epoch surrounding wait zone entry (- 0.5 to 0.5 s at wait zone entry). Normalized activity at WZ entry was averaged separately based on whether the encounter was quit, skipped or earned. Within each decision, activity was further subdivided into delays. To reduce noise resulting from sparse sampling of quit decisions that were further reduced by separating data by delay, we averaged across 11-20 s delays to define “medium delays” and 21-30 s delays to define “long delays” for each decision. A One-Way ANOVA was used to assess statistical differences across decisions.

*Figure 6 Reward Activity Analyses:* Normalized firing rate was averaged across cells within each subregion during the epoch surrounding reward delivery. We explored neural dynamics during the -3 to +3 s window around the time at which the computer turned on the feeder (t = 0 s refers to the moment the feeder clicked, indicating its operation). The “reward anticipation” period was defined as the -3-0 s period preceding the feeder click. It took approximately 1 s for food pellets to reach the receptacle on the maze where they were available for consumption. We therefore defined the 0-1 epoch following feeder click as the “reward delivery” period, referring to the collection of sounds that happened with incoming pellets. Given how quickly rats consumed food upon its arrival into the feeding receptacle, we defined the 2-3 s epoch following feeder click as the “reward consumption” period. A mixed effects model (2 fixed terms and 1 random term) was used to test for region-dependent differences in reward activity: fr ∼ region*epoch + (1|ratID).

Changes in reward-related activity with global economic conditions were assessed by measuring normalized firing rate averaged across cells within each subregion within each reward epoch (with temporal windows defined as above). A mixed effects model (2 fixed effects and 1 random effect) was used to test for effects of epoch and task stage on neural activity within each subregion: fr ∼ epoch*stage + (1|ratID).

Changes in reward-related activity with local economic conditions were assessed by measuring cell-averaged normalized firing rate binned in successive 5-min intervals within each session. Cell-averaged firing rates were then averaged across the temporal windows that defined each reward epoch. A mixed effects model (2 fixed effects and 1 random effect) was used to test for effects of session time on reward-related activity in each subregion: fr ∼ epoch*chunk + (1|ratID).

To examine how neural activity during the reward epoch was influenced by delay, we separated each reward delivery by the delay preceding its delivery. For each cell, we found its average normalized firing rate separately within reward anticipation, delivery, and consumption epochs for each delay. This resulted in 3 separate nCells×delays matrices, one for each reward epoch. We then averaged across cells within each subregion to obtain tuning curves of normalized firing rate to delay during reward anticipation, delivery, and consumption. A mixed effects model (2 fixed terms and 1 random term) was used to test for effects of delay and epoch on neural activity: fr ∼ delay*epoch + (1|ratID).

To explore representations of reward-specific attributes, we used a Bayesian decoding algorithm to estimate a posterior probability distribution over reward types during the reward epoch. This decoding algorithm takes as input a cells × session-time firing rate matrix along with a reward-type × cells matrix of tuning curves. Given the ensemble tuning curves, the algorithm uses ensemble firing rates at each moment, t, to estimate a probability distribution over reward types (flavors) at that moment. We constructed tuning curves to reward types during the 0-2 s epoch following reward delivery, consistent with previous research exploring reward representations in Restaurant Row. We cross-validated all decoding analyses by randomly selecting half of the encounters within a session to construct tuning curves (i.e., train the model) and analyzed decoded probability distributions from the other half of the session (i.e., test the model). We repeated this cross-validation 50 times per session and averaged across them. The bin size for constructing the cells × session time firing rate matrix, along with the peri-event bin size, was 250 ms. This resulted in a sessions × time × reward-type matrix of posterior probability distributions for each subregion within the OFC.

To analyze reward-specific representations, we rotated the maze so that the posterior probability distribution over reward types were aligned to the current, next, past, and other restaurant relative to the rat’s current position. For example, if the rat was currently at the restaurant offering cherry pellets, then the “current” reward would be assigned the probability of decoding to cherry. The “next” reward would be assigned the probability of decoding to banana. The “other” reward would be assigned the probability of decoding to plain. The “past” reward would be assigned the probability of decoding to chocolate. For the next encounter, the current-next-past-other assignment would be shifted 90 degrees on the maze, such that now the “current” reward was assigned the probability of decoding to banana. Thus, reward-specific representations emerged as a greater probability of decoding the current reward than the others.

To measure the strength of current reward representations during reward anticipation, delivery, and consumption, we averaged the posterior probability of each reward type across time separately within each epoch and found the difference between the probability assigned to current reward from the probability assigned to each of the other three. Thus, for each lap we obtained 3 values measuring the difference between current reward probability and next, other, and past reward probabilities. Values greater than 0 indicate that the current reward was more strongly represented than the other three. These values were averaged across each pairwise comparison and finally across laps to obtain, for each subregion within each session, a measure of the strength of reward-specific representations during each epoch surrounding reward.

As a control, we randomly assigned the rat’s current restaurant with one of the three remaining labels (i.e., next, other, past). The order (or structure) of the relation between current-next-past-other was preserved for each random assignment, which resulted in a more stringent control test because information about the order of reward flavors was preserved. In other words, the shuffle related to where within a fixed sequence of reward identities the rat was assumed to be at, rather than a complete shuffling of reward types and their spatial positions relative to the rat. As with the experimental data, we then averaged laps and across pairwise comparisons of the difference between the probability of the reward labeled as “current” with the other three reward positions. Significant current-reward encoding was determined using 1) Wilcoxon Signed Rank Tests on normalized distributions of current-reward scores, and 2) Mann-Whitney U Tests between real and shuffled distributions of current-reward scores.

*Figure 7: Distinct Neural Dynamics in LO and VO During Offer and Wait Zone Decisions*. Each cell’s average normalized firing rate during an epoch of interest was used to determine the degree to which that cell was sensitive to the behavioral event defining that epoch. For LO cells, we were interested in sensitivity to offer zone entry, wait zone entry, and reward delivery (linger zone entry). Normalized firing rate for each LO cell was averaged across a -0.5 to 0.5 s window at offer zone entry, a 0.5 to 1 s window at wait zone entry, and a 0-1 s window at reward delivery. For VO cells, we were interested in offer zone exit given skip, wait zone exit given quit, and linger zone exit. Normalized firing rate for each VO cell was averaged during the -2.5 to 0.5 s window prior leaving each of the three zones. Temporal windows used for averaging neural activity were determined based on ensemble-level neural dynamics at each behavioral epoch. A cell was deemed sensitive to an event if its normalized firing rate exceeded 1 standard deviation of the distribution mean of cell scores at that event. This method thereby selected those cells within the population that were most strongly active during the behavioral epoch of interest.

The correlation of normalized firing rate scores between each pair of behavioral epochs (offer zone entry, wait zone entry, and reward) was measured to determine the degree to which a cell’s activity at one epoch was related to its activity at another. In addition, we identified cells that were deemed sensitive to a behavioral epoch and plotted peri-event averages of activity within those cells at each behavioral epoch. In doing so, we were able to compare how the temporal dynamics of a collection of cells deemed sensitive to an event unfolded across behavioral epochs.

To examine the extent to which each cell’s tuning curve to momentary delay was consistent between the offer zone and wait zone, we normalized the correlation between tuning curves derived from each zone relative to a shuffled distribution. Specifically, for each cell, we preserved the relation between firing rate and momentary delay for a given zone (e.g., offer zone) and then randomly shuffled the relation between firing rate and momentary delay for the other zone (e.g., wait zone). We then found the correlation between these tuning curves. This process was repeated 100 times. The true correlation between tuning curves was then z-scored relative to the n = 100 shuffled distributions.

## Supplementary Information Figures

**S1.**
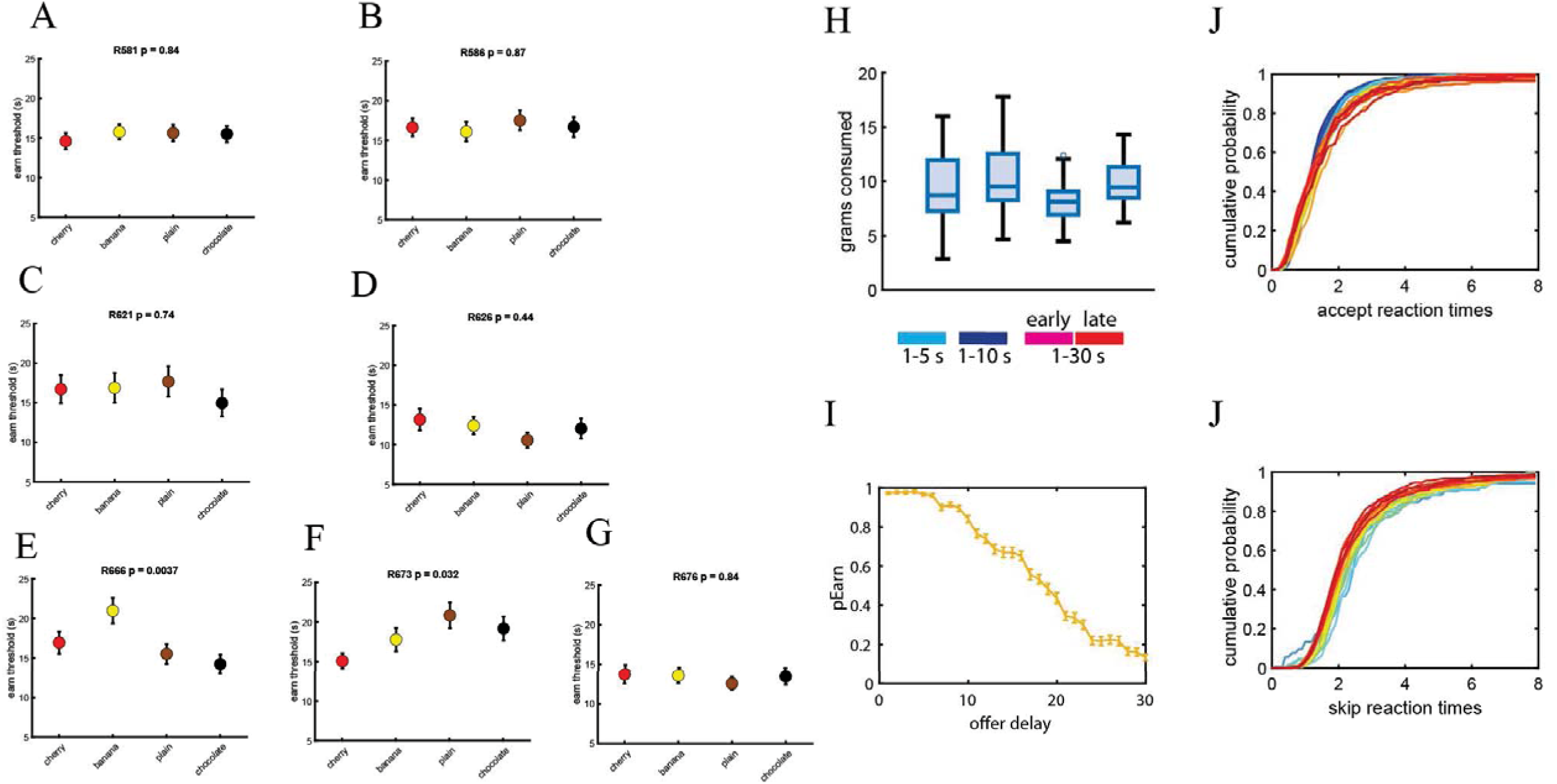
Behavior. (A-G) Mean (± SEM) earn threshold measured by fitting a heaviside function to a binary variable indicating whether reward was earned for each rat for each restaurants. The p values indicate the significance result from a mixed effects model testing for effects of restaurant on earn threshold. (H) Grams of food consumed across the four stages of training. (I) Mean (± SEM) probability of earning food as a function of delay during the reward scarce environment. (J-K) Probability density functions of time spent in the OZ as a function of delay during accepts (J) and skips (k). Reaction times for each delay are broken up by color, with redder colors indicating longer delays.

**S2.**
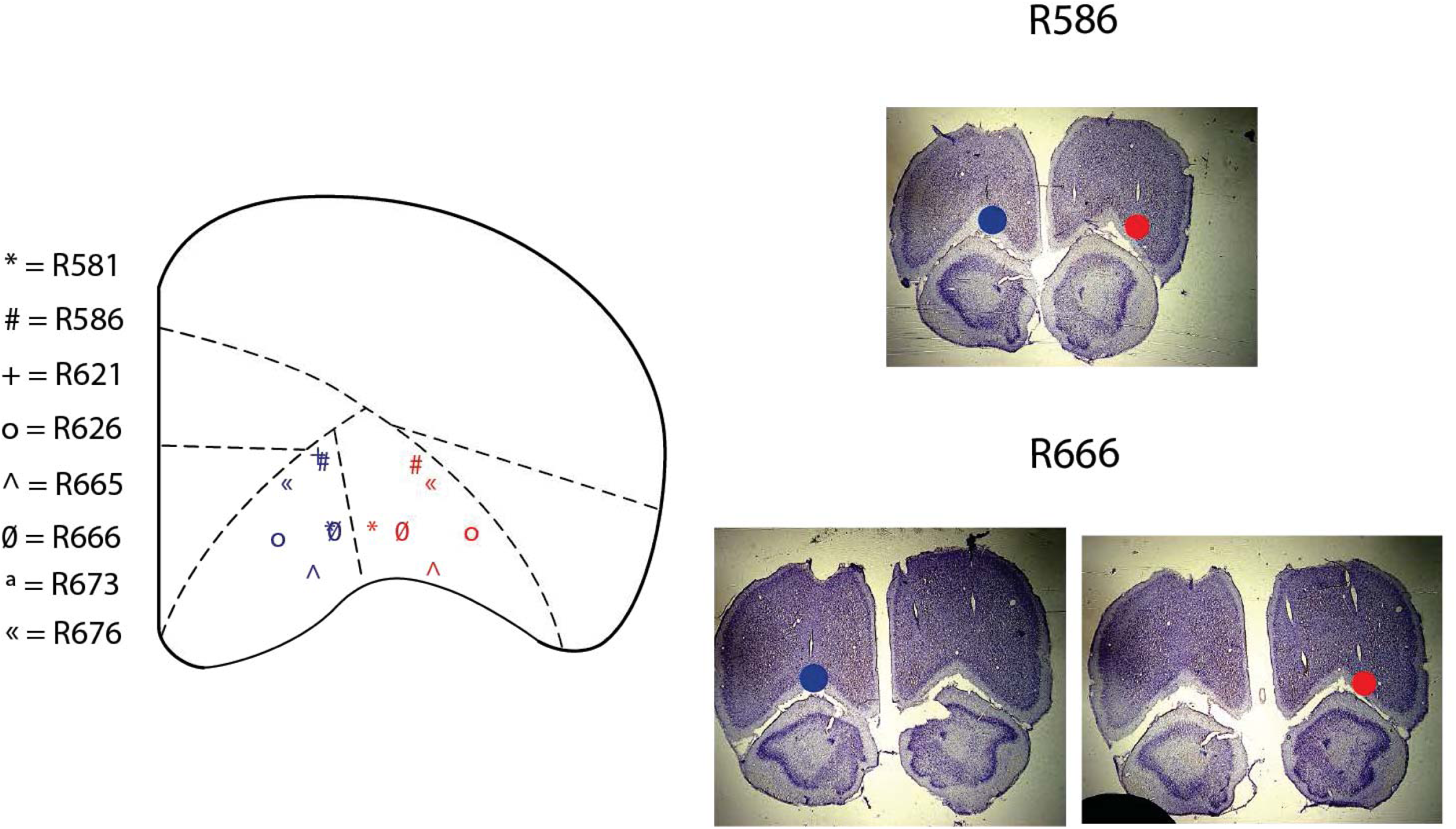
Histology. Probe tracks for LO and VO targets are depicted across rats, along with images from two example rats.

**S3.**
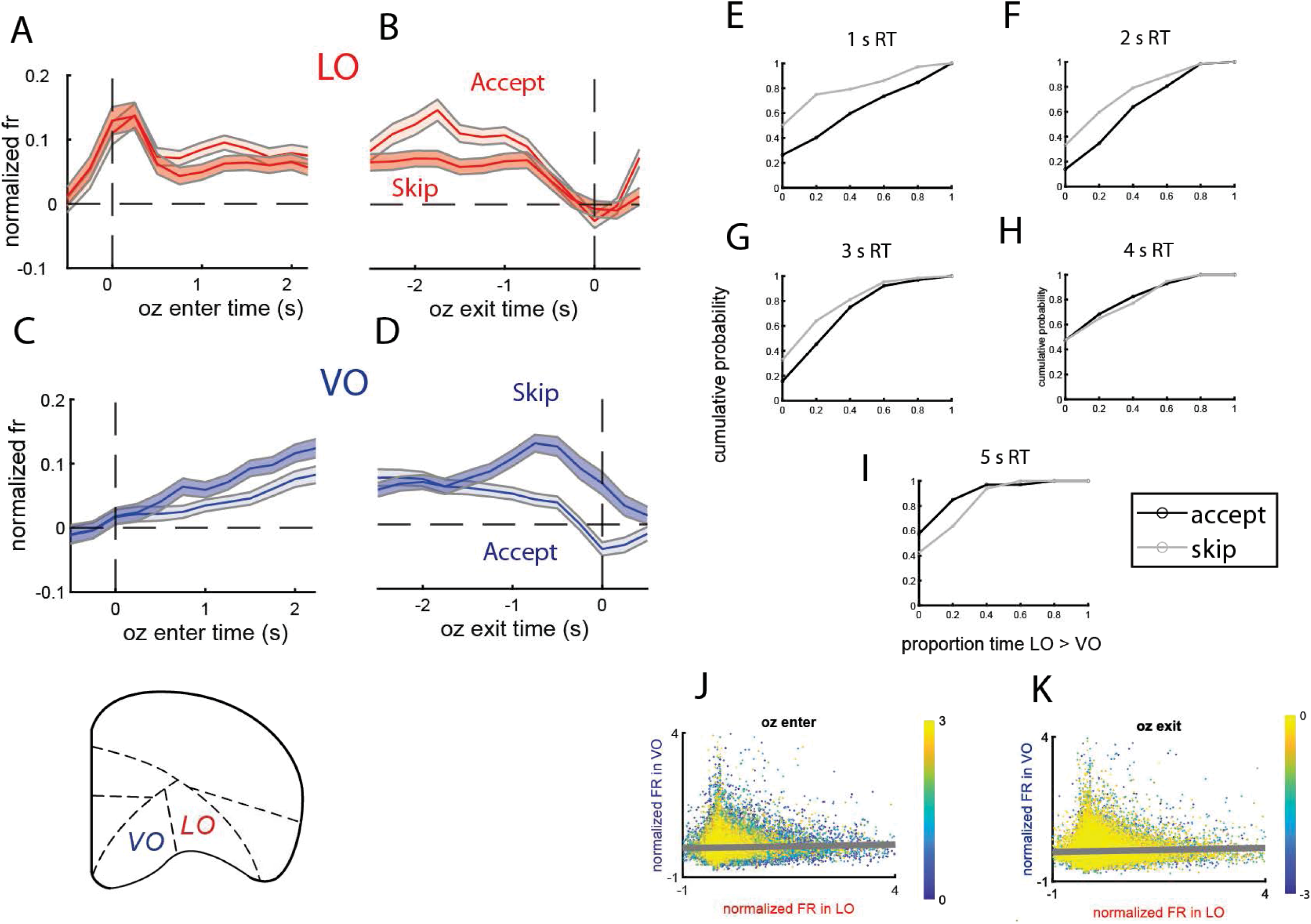
Decision-related activity in the offer zone across subregions. (A-D) Mean (± SEM) normalized firing rates during accept and skip encounters in LO (A-B), VO (C-D). The left panels show normalized activity oriented with respect to offer zone enter time while the right panels show normalized activity at offer zone exit time. (E-I) Cumulative density functions of the proportion of time in the offer zone in which LO activity exceeded VO activity for accept and skip encounters. Each panel shows these cumulative density functions broken up by the amount of time (reaction time – RT) spent in the offer zone. (J-K) Scatter plot of the moment-by-moment correlation of LO and VO ensemble activity across all sessions during the first and last 3 s of OZ entry (J) and OZ exit (K). The colormap indicates the progression of time, and each point shows average activity in LO (x axis) and VO (y axis) ensembles for a given session.

**S4.**
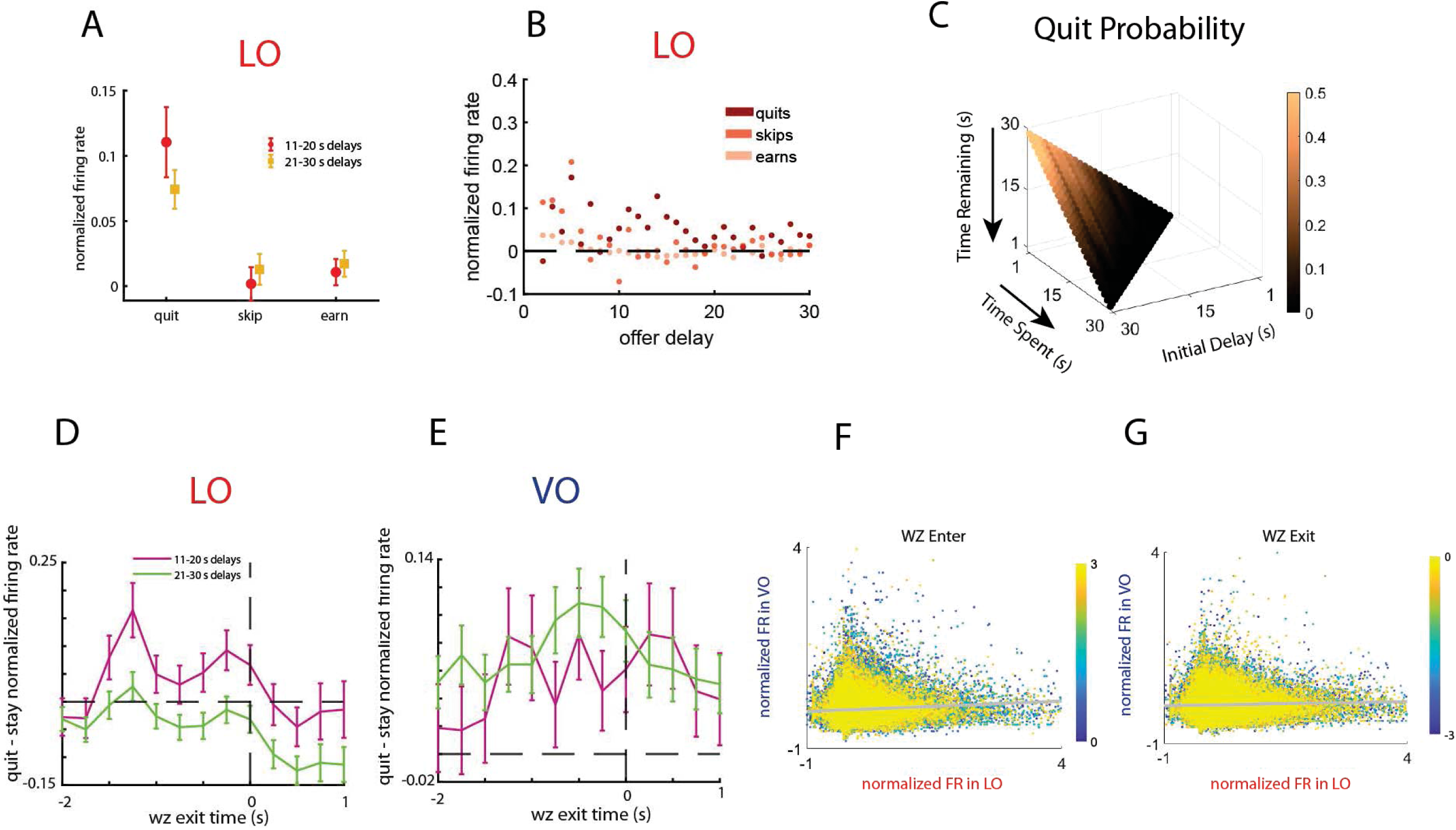
Decision-related activity in the wait zone across subregions. (A) To explore the possibility that heightened LO activity at wait zone entry during quits resulted from abnormally high LO activity for long-delay offers in the offer zone, resulting in an erroneous accept decision, we compared LO activity for quits, skips, and earns separately for medium and long delays. Mean (± SEM) LO activity during the epoch surrounding offer zone exit for encounters that were quit, skipped, or earned during medium and long delay offers. (B) Tuning curves of normalized LO firing rate to offer delay during the epoch surrounding offer zone exit when the offer was skipped, earned, and quit. (C) Quit probability as a function of time remaining and time spent across offer delays. (D-E) Mean (± SEM) difference in neural activity between quit and stay encounters separately for medium (11-20 s) and long (21-30 s) delays. Mean (± SEM) difference in quit from stay normalized firing rate, oriented with respect to wait zone exit, in the LO (C) and VO (D) averaged across medium and long delays. (F-G) Scatter plot of the moment-by-moment correlation of LO and VO ensemble activity across all sessions during the first and last 3 s of WZ entry (F) and WZ exit (G). The colormap indicates the progression of time, and each point shows average activity in LO (x axis) and VO (y axis) ensembles for a given session.

**S5.**
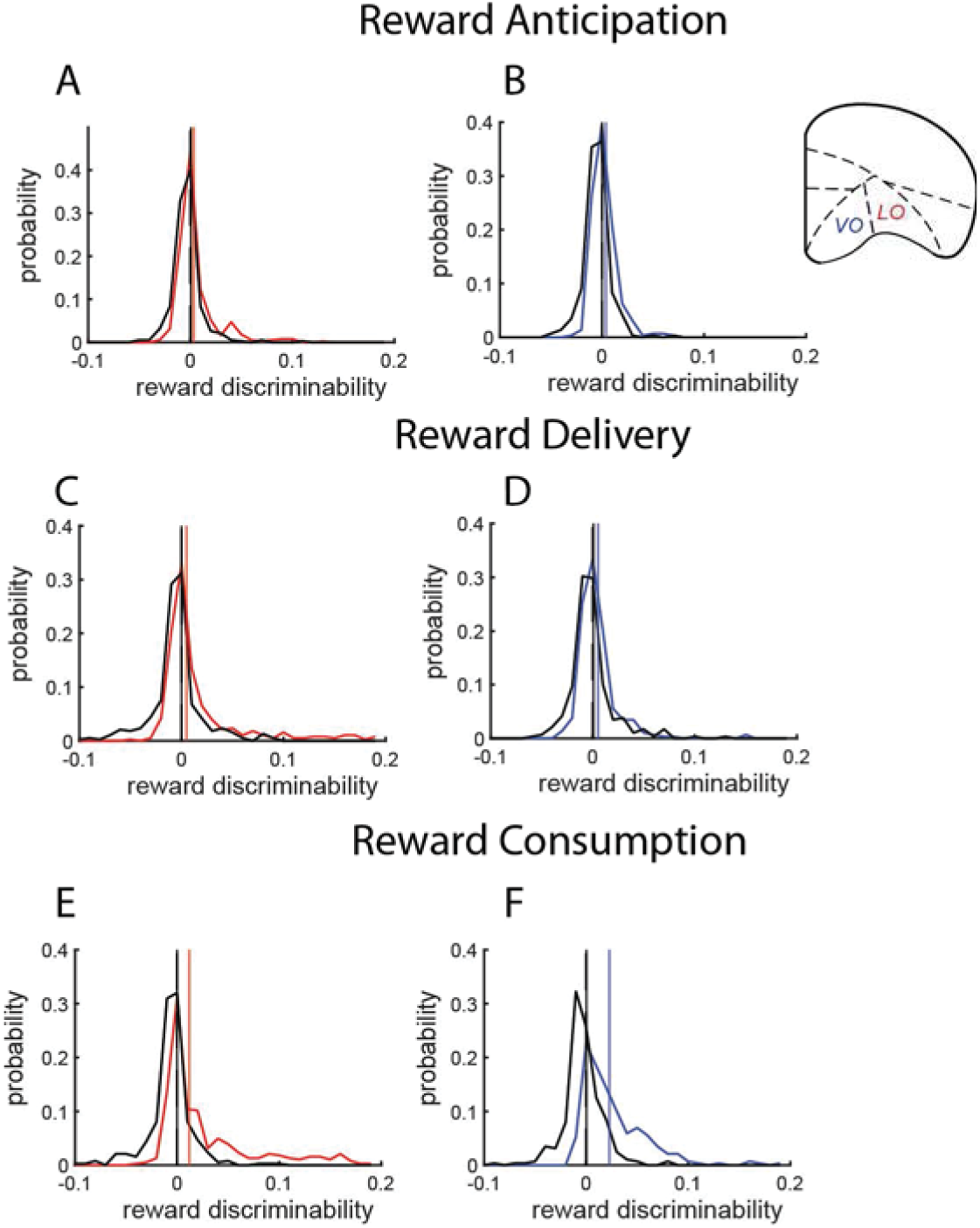
Distributions of reward-specific representations across the OFC during anticipation, delivery, and consumption. To quantify the degree to which ensemble activity encoded specific features of reward, we measured the mean difference in the probability of decoding to the current reward relative to each of the other three rewards. We compared the distribution of these scores to a shuffled distribution which randomly assigned reward flavors to current, next, past, and other rewards (see Methods for details). Positive values indicate information about specific rewards, and more positive values indicate a greater degree of information about current reward. The distribution of current reward decoding scores during reward anticipation (A-B), delivery (C-D), and consumption (E-F) are depicted for LO (left panels) and VO (right panels). Shuffled distributions are in black. Vertical lines reflect medians of the real and shuffled distributions. Stats performed on these distributions are reported in the main text.

## Supplementary Information Tables

**Table S1:** Summary of recording information and cell counts across rats.

|  | Sex | Probe Type (# of channels) | Subregion | Hemisphere | Cell Count |
| --- | --- | --- | --- | --- | --- |
| **R581** | F | F-probe (64) | LO<br>VO | Right<br>Both | 213<br>567 |
| **R586** | M | E2-probe (32) | LO<br>VO | Left<br>Right | 214<br>593 |
| **R621** | F | E2-probe (32) | LO<br>VO | Left<br>Right | 132<br>164 |
| **R626** | F | E2-probe (32) | LO<br>VO | Left<br>Right | 60<br>115 |
| **R665** | M | E2-probe (32) | LO<br>VO | Right<br>Left | 0<br>0 |
| **R666** | M | E2-probe (32) | LO<br>VO | Left<br>Right | 31<br>787 |
| **R673** | M | E2-probe (32) | LO<br>VO | Right<br>Left | 667<br>0 |
| **R676** | F | E2-probe (32) | LO<br>VO | Right<br>Left | 111<br>0 |

